# The hologenome of *Osedax frankpressi* reveals the genetic interplay for the symbiotic digestion of vertebrate bone

**DOI:** 10.1101/2022.08.04.502725

**Authors:** Giacomo Moggioli, Balig Panossian, Yanan Sun, Daniel Thiel, Francisco M. Martín-Zamora, Martin Tran, Alexander M. Clifford, Shana K. Goffredi, Nadezhda Rimskaya-Korsakova, Gáspár Jékelly, Martin Tresguerres, Pei-Yuan Qian, Jian-Wen Qiu, Greg W. Rouse, Lee M. Henry, José M. Martín-Durán

## Abstract

The marine annelid *Osedax* has evolved a unique heterotrophic symbiosis that allows it to feed exclusively on sunken bones. Yet, the genetic and physiological principles sustaining this symbiosis are poorly understood. Here we show that *Osedax frankpressi* has a small, AT-rich genome shaped by extensive gene loss. While the *Oceanospirillales* endosymbiont of *Osedax* is enriched in genes for carbohydrate and nitrogen metabolism, *O. frankpressi* has undergone genetic changes to accommodate bone digestion, including the expansion of matrix metalloproteases, and a loss of pathways to synthesize amino acids that are abundant in collagen. Unlike other symbioses, however, innate immunity genes required to acquire and control the endosymbionts are reduced in *O. frankpressi*. These findings reveal *Osedax* has evolved an alternative genomic toolkit to bacterial symbiosis where host-symbiont co-dependence has favoured genome simplicity in the host to exploit the nutritionally unbalanced diet of bones.

**Teaser:** Genome reduction and adaptations for collagen digestion underpin the symbiosis of *Osedax* worms to exploit decaying bones.

## Introduction

Symbioses have shaped life on Earth, from the origin of the eukaryotic cell to the formation of biodiversity hotspots such as coral reefs (*1, 2*). Animal chemosynthetic symbioses, where bacteria convert inorganic compounds to organic matter, are ubiquitous in marine habitats (*3*) and fuel some of the most productive communities, such as those around hydrothermal vents (*4*). Siboglinid worms (Annelida) often dominate deep-sea chemosynthetic environments through symbioses with environmentally acquired bacteria (*5, 6*) that adults harbour within a specialised organ called a trophosome (*7*). Despite their ecological importance, our understanding of the genetic traits that sustain symbioses in Siboglinidae is scarce and currently limited to Vestimentifera (*8–10*), one of the four main lineages in this annelid group (Figure 1A). Loss of genes involved in amino acid biosynthesis (*8, 10*) and carbohydrate catabolism (*9*) together with expansions of genes involved in nutrient transport (*8*), gas exchange (*8–12*), innate immunity and lysosomal digestion (*8–10, 13*) show a complex molecular interplay between Vestimentifera and their endosymbionts to fulfil their nutritional demands (*14*). Notably, many of these genetic changes are common to Vestimentifera dwelling in both methane seeps and hydrothermal vents, and also to other distantly related chemosymbiotic animals such as bivalves (*15*), gastropods (*16*), and the clitellate annelid *Olavius algarvensis* (*17*). Therefore, disparate groups of animals have convergently evolved distinct genetic mechanisms that underpin the emergence of chemosynthetic symbioses in marine ecosystems.

**Figure 1.**
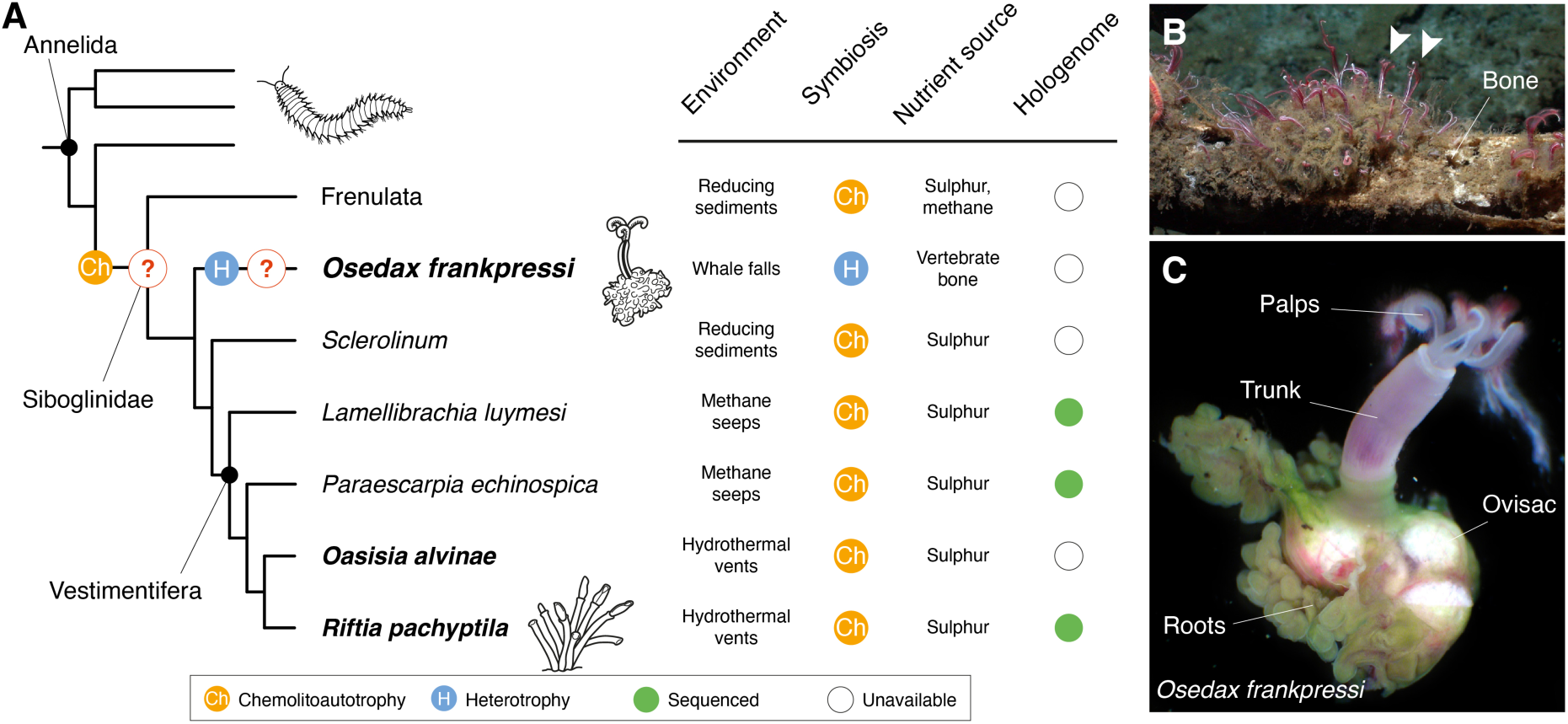
Siboglinidae is a symbiotic annelid group. (**A**) Siboglinidae is a diverse clade of annelid worms that evolved chemosynthetic symbioses (left side). There are four main lineages within Siboglinidae, namely Frenulata, *Osedax*, *Sclerolinum* and Vestimentifera. Chemolithoautotrophy occurs in Frenulata, *Sclerolinum* and Vestimentifera, which associate with gammaproteobacteria that employ sulphur or methane to produce organic compounds in an array of marine ecosystems, from reducing sediments to methane seeps and hydrothermal vents (right side of the panel). Differently, *Osedax* worms (e.g., *O. frankpressi*; **B** and **C**) have secondarily evolved a heterotrophic association with *Oceanospirillales* to exploit decaying vertebrate bones. The genomic basis for the evolution of these nutritional symbioses in Siboglinidae is unclear (question marks on the left) because genomic information only exists for three Vestimentifera species (green circles on the right). The species herein studied are highlighted in boldface. (**B**, **C**) Photographs of *O. frankpressi* in a whale bone (**B**; arrowheads point to *O. frankpressi*) and a mature female adult (**C**). *O. frankpressi* settles and colonises decaying vertebrate bones (**B**). There, the posterior part of the body becomes stably infected with environmentally acquired *Oceanospirillales* bacteria. This body part (the so-called roots) harbours the bacteria and grows to penetrate the bone, dissolving the organic components. These nutrients are absorbed and transported towards the bacteriocytes containing the endosymbionts, which will proliferate and act as food for the worm. Anterior to the root tissue there are the reproductive ovisacs and the head bears two pairs of palps.

Within Siboglinidae, the marine bone-eating annelids *Osedax* have evolved a remarkable symbiosis that is unique among animals (*18–22*) (Figure 1A). *Osedax* consumes bones aided by endosymbiotic heterotrophic bacteria in the order *Oceanospirillales* (*19, 23–26*) (Figure 1B). While *Osedax* shares some morphological features with other siboglinids (*27*), including the lack of a gut, mouth and anus, *Osedax* contains bacteriocytes that are concentrated in the subepidermal connective tissue of the lower trunk that grows directly into the bone (Figure 1C) (*19, 23*). This amorphous tissue is referred to as “roots” and expresses high levels of V-type H^+^-ATPase and carbonic anhydrase, indicating acid is used to dissolve the bone matrix to access collagen and lipids (*28*), which are then absorbed across the root epithelium. Enzymatic (*23, 25*) and transcriptomic data (*29*) support this theory by showing the roots of *Osedax* express a large number of proteases and solute carrier transporters that are thought to be involved in bone degradation and nutrient absorption, perhaps with the aid of the endosymbionts (*24*). However, it is currently unclear whether the specialized heterotrophic symbiosis of *Osedax* is based on homologous genetic traits to those discovered in Vestimentifera, or if it relies on unique genomic adaptation to its unusual lifestyle. Untangling the molecular mechanisms behind the symbiosis of *Osedax* is therefore central to understanding the ecological principles and succession of bone-eating communities (*30*).

In this study, we sequenced the hologenome of *Osedax frankpressi* Rouse, Goffredi & Vrijenhoek, 2004, as well as that of two vent dwelling Vestimentifera, *Oasisia alvinae* Jones, 1985 and *Riftia pachyptila* Jones, 1981, for comparative purposes. In contrast to Vestimentifera, we found that *O. frankpressi* has a small AT-rich genome with a reduced gene repertoire. Gene families that are typically expanded in chemosymbiotic hosts, such as innate immunity components, are highly reduced in *O. frankpressi*, which instead has unique genomic adaptations for bone digestion, including the loss of biosynthetic pathways of amino acids that are abundant in vertebrate bones and expansions of matrix metalloproteases that are important in bone digestion. Together, our findings demonstrate that divergent genomic adaptations sustain the nutritional symbioses of *Osedax* and Vestimentifera, providing key insight into the genetic and metabolic adaptations that have enabled symbiotic siboglinids to colonize diverse nutrient imbalanced feeding niches in the sea.

## Results and Discussion

### *The hologenomes of* O. frankpressi, Oasisia alvinae *and* R. pachyptila

To identify genomic signatures that could inform the genetic and physiological basis of the heterotrophic symbiosis in *Osedax*, we used long PacBio reads and short Illumina reads to assemble the genome of *O. frankpressi* (*19*) (Figure 1B, C; Supplementary Table 1). We also sequenced the genomes of two Vestimentifera from hydrothermal vents, *Oasisia alvinae* and *R. pachyptila* (Supplementary Figure 1), which complement previous genome sequencing efforts (*8–10*). We generated almost completely de-haploidised draft assemblies (Supplementary Figure 2A–D), which included the circularised endosymbiont genomes of *O. frankpressi* and *Oasisia alvinae* and several epibionts associated with *O. frankpressi* (Supplementary Figure 2H–J; Supplementary Table 2). Consistent with *k*-mer-based analyses (Supplementary Figure 2E–G), previously reported genome size estimation for *Oasisia alvinae* (*31*), and a recent genome assembly of *R. pachyptila* (*10*), the assembled genomes for *O. frankpressi*, *Oasisia alvinae* and *R. pachyptila* span 285 Mb, 808 Mb and 554 Mb after removal of bacterial contigs, respectively (Figure 2A; Supplementary Figure 2K). For *Oasisia alvinae* and *R. pachyptila*, the genome assembly shows high completeness (96.9% and 95.6% BUSCO presence, respectively; Supplementary Figure 2L; Supplementary Table 3), yet the assembly for *O. frankpressi* appeared to have a lower completeness (80.1% BUSCO presence; Supplementary Figure 2L). However, 95.62% and 97.77% of the *de novo* assembled transcripts from the body and root tissue mapped to the genome assembly of *O. frankpressi*, respectively, and thus BUSCO completeness increased to a final score of 96.23% after gene annotation (Supplementary Figure 2L) and manual curation (26 out of the 62 missing BUSCO could be manually annotated; Supplementary Table 4). The relatively low assembly-based BUSCO completeness is therefore compatible with the fast rates of molecular evolution in coding sequences observed in *Osedax* worms (*32*).

**Figure 2.**
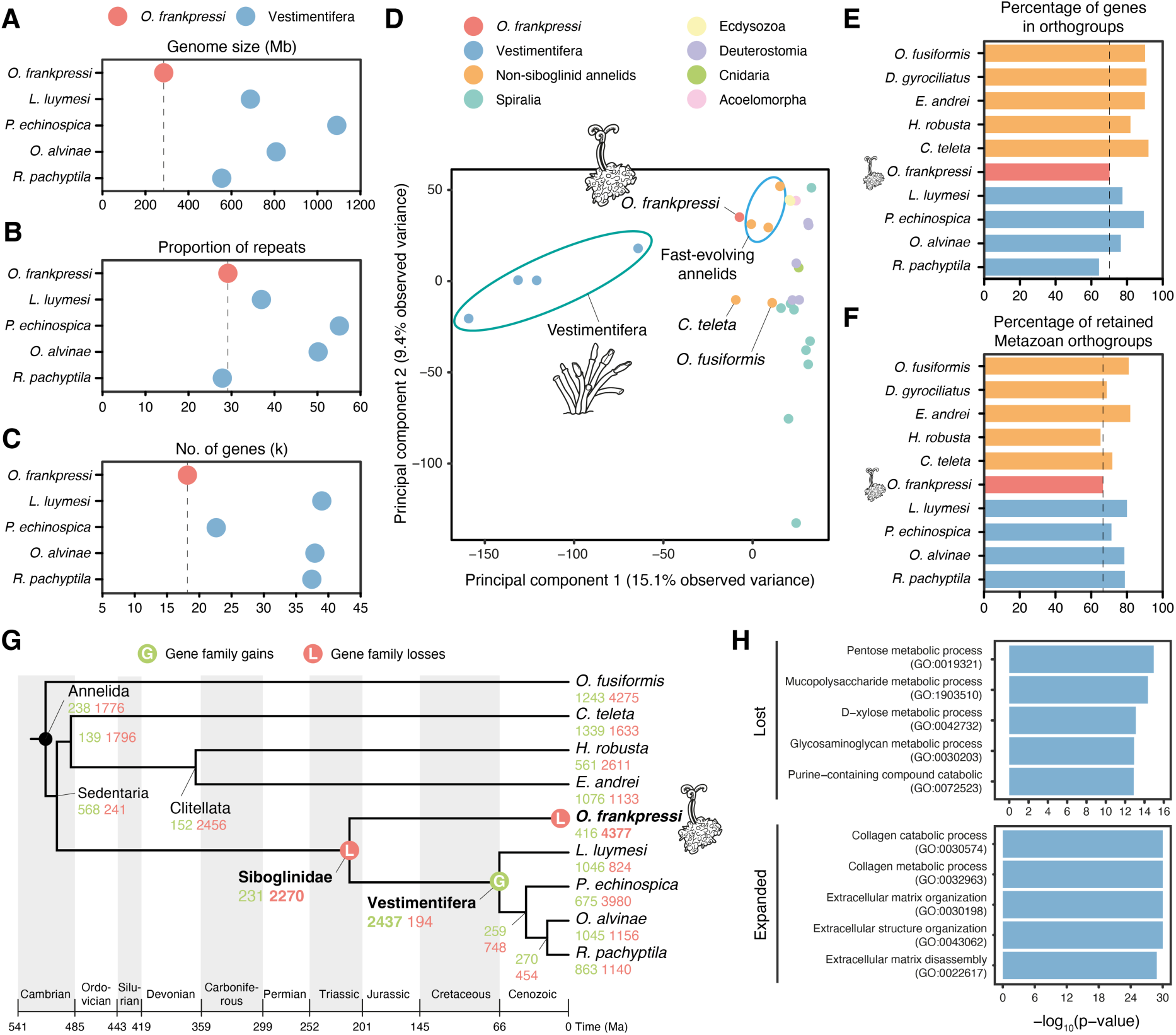
*Osedax frankpressi* has a small genome and a reduced gene repertoire. (**A**–**C**) Plots comparing genome size (**A**), repeat content (**B**) and number of genes (**C**) between *O. frankpressi* and the four Vestimentifera with sequenced genomes. *O. frankpressi* has a smaller genome, with less genes but relatively similar repeat content. (**D**) Principal component analyses of the gene content of 28 metazoan genomes show that differently from symbiotic bivalves and gastropods, the gene content of Vestimentifera and *O. frankpressi* differs from slow-evolving asymbiotic species (as represented by *Owenia fusiformis* and *C. teleta*). While Vestimentifera has a unique gene content, *O. frankpressi* is like other fast-evolving annelid lineages. (**E**, **F**) Bar plots of the percentage of genes in gene families (i.e., orthogroups; **E**) and retained ancestral metazoan gene families (**F**) for ten annelid lineages. *O. frankpressi* is amongst the annelids with less genes in gene families and less retained ancestral metazoan genes. (**G**) Patterns of gene family gains (in green) and loss (in red) during the evolution of Annelida under a consensus tree topology (*27*) and a consensus of published molecular dates (*8, 9*). A major event of gene loss is common to all Siboglinidae. While *O. frankpressi* continued experiencing high rates of gene loss, a major event of gene innovation is common to all Vestimentifera. (**H**) Top five enriched gene ontology terms (Biological Process) for gene families lost (top) and expanded (bottom) in *O. frankpressi*. While *O. frankpressi* has further lost genes involved in metabolism (e.g., carbohydrate metabolism), genes involved in collagen and extracellular matrix degradation are expanded.

Although the genome of *O. frankpressi* is ∼50–75% smaller than the sequenced genomes and estimated genome sizes of Vestimentifera (Figure 2A) (*8–10, 31*), the fraction of simple repeats and transposable elements in *O. frankpressi* (29.16%) is comparable to that of the vestimentiferan *R. pachyptila* (27.87%) and asymbiotic annelids with similar genome sizes, such as *Capitella teleta* (31%) (Figure 2B; Supplementary Figure 3A). Moreover, as in Vestimentifera, but unlike asymbiotic annelids with slow rates of molecular evolution such as *Owenia fusiformis* and *C. teleta* (*33, 34*), the repeat landscape in *O. frankpressi* shows signs of expansions (Supplementary Figure 3B). Combining transcriptomic evidence (Supplementary Table 1) with *ab initio* gene prediction (Supplementary Figure 2A), we functionally annotated 37,929 and 37,455 protein coding genes in *Oasisia alvinae* and *R. pachyptila*, respectively (Supplementary Figure 2H), which have similar number of genes to other Vestimentifera and asymbiotic annelids (*8, 33, 34*). In contrast, *O. frankpressi* has a smaller repertoire of 18,808 genes (Figure 2C), comparable to that of the miniaturised *Dimorphilus gyrociliatus*, another annelid species with a compact genome and a streamlined gene set (14,203 genes) (*32*). Therefore, *O. frankpressi* has the smallest genome of all sequenced Vestimentifera. Given the number of genes in genomes of asymbiotic annelids, gene loss rather than removal of repeat content seems to account for the genome size difference between these two lineages of Siboglinidae.

### Gene family gains and losses shape the evolution of Siboglinidae

To investigate gene content evolution between major lineages of Siboglinidae, we first conducted principal component analysis of 28 highly complete metazoan genomes, including seven symbiotic annelid and molluscan lineages (Supplementary Table 8). The symbiotic molluscs *Bathymodiolus platifrons* (*15*) and *Gigantopelta aegis* (*16*) cluster with their asymbiotic bivalve and gastropod relatives, respectively (Supplementary Figure 4A). However, the four Vestimentifera species are markedly differentiated from the other annelid and animal genomes, and *O. frankpressi* is closer to heterotrophic annelids with fast rates of molecular evolution and divergent gene repertoires, such as the leech *Helobdella robusta* and the earthworm *Eisenia andrei*—which also harbour bacterial symbionts (*35–37*)—and the marine worm *D. gyrociliatus* (Figure 2D; Supplementary Figure 4A). Indeed, after *R. pachyptila*, *O. frankpressi* is the annelid with the second lowest percentage of genes assigned to gene families (Figure 2E) and has only retained a fraction of ancestral metazoan gene families comparable to more rapidly evolving annelids such as *H. robusta* and *D. gyrociliatus* (Figure 2F). Therefore, unlike symbiotic molluscs, the evolution of nutritional symbioses in Siboglinidae correlates with divergent host gene repertoires compared to their asymbiotic annelid counterparts.

To identify and characterise the evolutionary events underpinning the divergent gene repertoires of Siboglinidae, we reconstructed the patterns of gene family evolution in those 28 metazoan genomes under a consensus tree topology (Supplementary Figure 4B). A major event of gene loss involving 2,270 gene families of mostly ancient origins (61.23% of the lost families originated prior to Metazoa and/or the Bilateria/Nephrozoa ancestor) is common to Vestimentifera and *O. frankpressi* (Figure 2G) and largely involves genes enriched in Gene Ontology (GO) terms associated with metabolism (Supplementary Figure 4C). This loss thus coincides with the evolution of nutritional symbioses in the last common ancestor of Siboglinidae. High rates of gene loss continued in the *O. frankpressi* lineage (Figure 2G), which ultimately account for its reduced gene repertoire and largely affected genes associated with carbohydrate and nitrogen metabolism (Figure 2H; Supplementary Figure 4D). Conversely, Vestimentifera experienced an event of gene family expansion in its last common ancestor (2,437 gene families), mostly affecting genes related to immunity, cell communication, and response to stimuli (Figure 2G; Supplementary Figure 4E) (*9*). In contrast, *O. frankpressi* has had few gene family gains (Supplementary Figure 4B) but has experienced a large expansion of gene families associated with extracellular matrix remodelling and degradation (e.g., collagen degrading proteases; Figure 2H; Supplementary Figure 4F), in agreement with previous transcriptomic observations (*29*). Altogether, our findings indicate that the evolution of symbiosis in *Osedax* and Vestimentifera relies on markedly different host gene repertoires, one sculptured predominantly through gene loss (in *O. frankpressi*) and another through gene gains (in Vestimentifera) (*8–10*) (Figure 2G).

### *Osedax* endosymbiont is enriched in metabolic capabilities

In animal nutritional symbioses, the microbial partner provides metabolic capabilities that meet dietary needs of the host (*3*), such as the production of organic matter in the nutrient-poor vents by the endosymbionts of Vestimentifera (*5*). To investigate the genetic and functional contribution of the endosymbionts to the nutritional symbiosis of *Osedax*, we used our PacBio long-read data to assemble the genomes of the primary endosymbionts of *O. frankpressi* (Rs1 ribotype) (Figure 3A), and *Oasisia alvinae* (Supplementary Figure 5), as well as several epibionts associated with *Osedax* (*38*). The circularised assembly of *Osedax*’s endosymbiont improved the previously published genome (*24*), revealing 95 new functional genes that provide additional insights into its symbiosis (Supplementary Table 9). Compared to deep-sea free-living relatives, the *Osedax* endosymbiont has a genome that is enriched in metabolic genes for protein secretion systems, carbohydrate metabolism, and coenzyme and amino acid biosynthesis (Supplementary Table 10B). This includes additional virulence factors, such as multiple complete copies of the Type 5a, 5b, and 6i secretion system pathways (Supplementary Table 10C) that are important for modulating interactions with other bacteria and eukaryotic hosts. *Neptunomonas japonica*, a close relative of the *Osedax* endosymbiont that was recovered from marine sediments near a whale fall, has many of the same metabolic capabilities of the endosymbiont; however, it lacks the additional secretion systems, which may explain why this bacteria does not establish symbiosis with *Osedax* (*39*).

**Figure 3.**
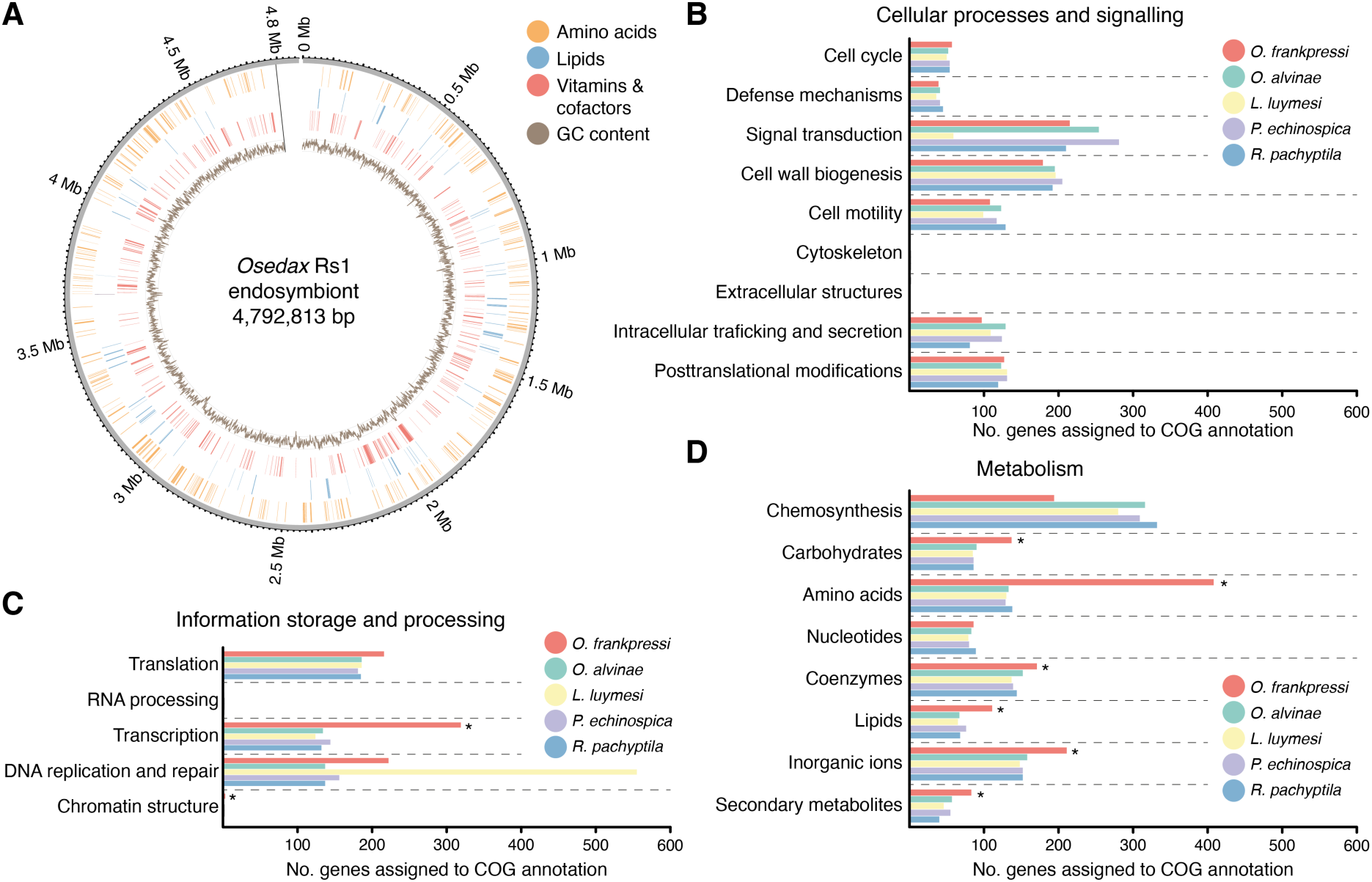
The endosymbiont of *O. frankpressi* has an enlarged metabolic repertoire. (**A**) Circular schematic representation of the genome of *Osedax* endosymbiont Rs1, assembled into a single contig. The plot shows the genomic location of genes involved in amino acid, lipid and vitamin/cofactor metabolism (in orange, blue and red, respectively) and the GC content (inner circle; brown colour). (**B**–**D**) Bar plots of number of COG annotations involved in cellular processes and signalling (**B**), information storage and processing (**C**), and metabolism (**D**) in the genome assemblies of *Osedax* endosymbiont and those of the four Vestimentifera with sequenced and assembled hologenomes. While there are no significant differences in the repertoire of genes involved in cellular processes and signalling between the endosymbionts of *Osedax* and Vestimentifera (**B**), *Osedax*’s endosymbiont has a significantly richer metabolic repertoire for carbohydrates, amino acids, coenzymes, lipids, inorganic ions, and secondary metabolites (**D**; significance indicated with an asterisk). Consistent with their chemosynthetic capacity, however, the endosymbionts of Vestimentifera are enriched in genes involved in energy production (**D**). The endosymbiont of *Osedax* is also enriched in genes involved in transcription and chromatin structure (**C**).

A comparison of *Osedax’s* endosymbiont to those of Vestimentifera reveals significant differences in their metabolic capabilities. While the repertoire of genes involved in core cellular processes was largely similar amongst all bacteria (Supplementary Table 11A), the *Oceanospirillales* endosymbiont has significantly more genes involved in the metabolism and transport of amino acids, coenzymes, lipids, and carbohydrates (Figure 3B–D; Supplementary Table 11A, E, F). This includes several complete pathways for the catabolism of glucose to pyruvate and multiple sugar transport system ATP-binding proteins (Supplementary Table 12). Most notably, the endosymbiont of *Osedax* can produce all essential amino acids, including methionine and threonine and vitamins B2, B6 and a complete B12 pathway, which chemolithoautotrophic endosymbionts from Vestimentifera cannot (Figure 4A; Supplementary Table 11D, E). Importantly, the B2 pathway was thought to be missing in the previous draft genome of the *Oceanospirillales* endosymbiont (*24*), but it is present in ours. As expected, all endosymbionts of Vestimentifera are enriched in genes involved in chemosynthesis, most of which are absent in the heterotroph endosymbiont of *Osedax* (Figure 3D).

**Figure 4.**
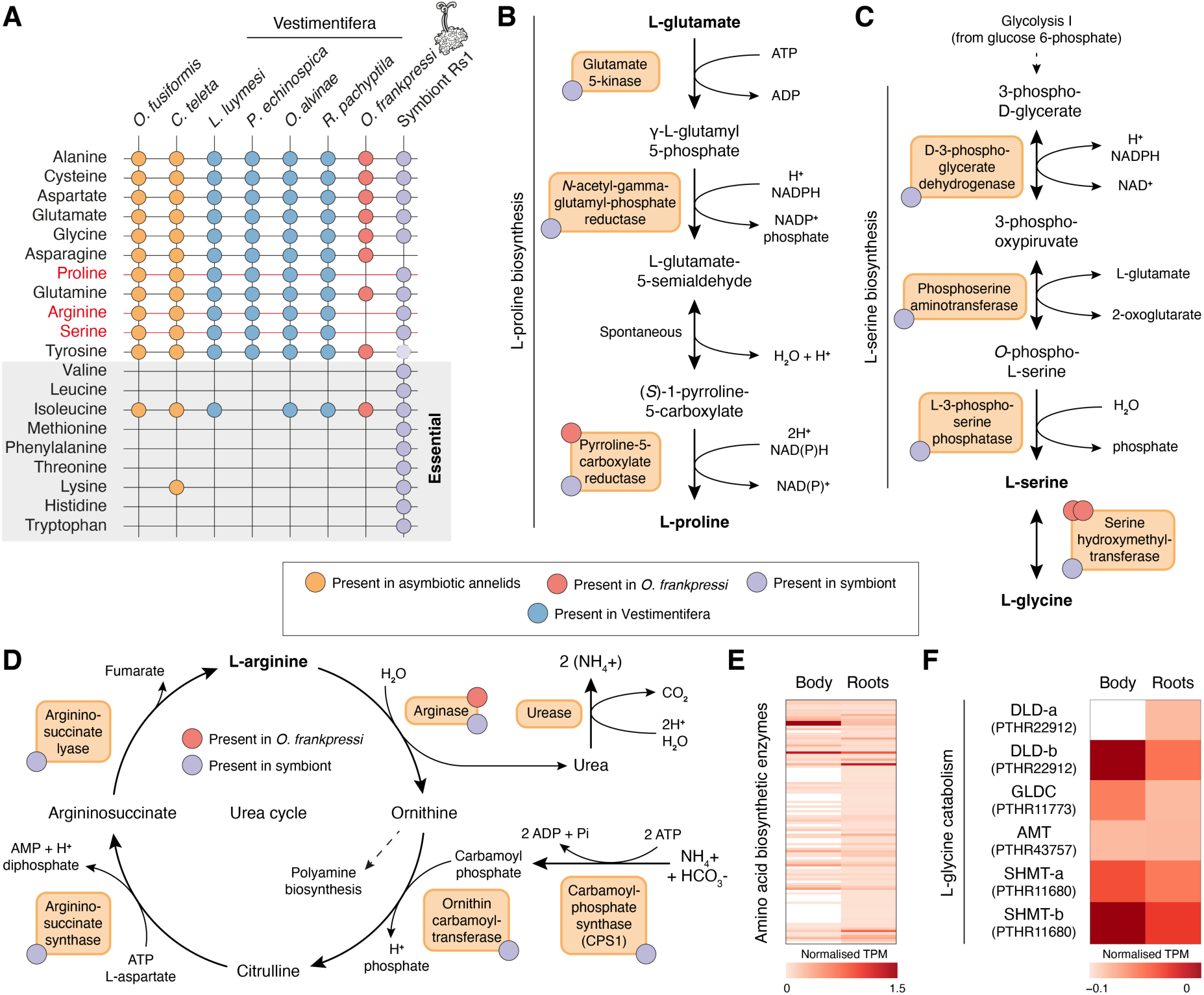
*O. frankpressi* shows metabolic adaptations to bone digestion. (**A**) Summary table of the presence (filled circles) and absence (empty crosses) of amino acid biosynthetic pathways in seven annelid genomes and *O. frankpressi* endosymbiont (symbiont Rs1). While Vestimentifera and asymbiotic annelids can synthesise all amino acids that are non-essential and conditional for humans, *O. frankpressi* shows incomplete pathways to synthetise proline, arginine, and serine (in red). Some of these amino acids are abundant in the bone (e.g., proline) and all can be produced by the symbiont (tyrosine biosynthetic pathway is truncated in the symbiont; dotted and lighter circle). (**B**–**D**) Schematic representation (as in MetaCyc database) of the biosynthetic pathways for proline (**B**), serine (**C**) and arginine (**D**) indicating with red and violet circles the enzymes present in *O. frankpressi* and its endsymbiont, respectively. *Osedax frankpressi* cannot produce serine from glycolytic metabolites but can either produce serine from collagen-derived glycine or take it from the diet. In addition, *O. frankpressi* can only convert arginine into ornithine, producing urea as a result. (**E**, **F**) Heatmaps of normalised mRNA expression levels for amino acid biosynthetic enzymes (**E**) and glycine catabolising enzymes (**F**) in the body and roots of *O. frankpressi*. Biosynthetic enzymes (**E**), including the two copies of serine hydroxymethyltransferase (SHMT-a and SHMT-b) that convert glycine into serine, are more expressed in the roots than in the body of *O. frankpressi*.

*Oceanospirillales* contains an abundance of secretion system that are absent in the symbionts of Vestimentifera. This includes three intact copies of both the Type 5a and Type 5b secretion systems (Supplementary Table 11G). This increase in virulence factors may reflect the fact that *Oceanospirillales* must repeatedly infect the roots of *Osedax* as it grows through bone material, unlike the trophosome of Vestimentifera that is colonised early during host development (*7, 40*). Taken together, our results further demonstrate that the heterotrophic endosymbiont of *Osedax* has a much more versatile metabolism than the chemolithoautotrophic bacteria of Vestimentifera.

### *Osedax* has metabolic adaptations to bone digestion

Vertebrate bones are nutrient imbalanced food sources (*41*). The reduced gene repertoire of *O. frankpressi* (Figure 2C) combined with the metabolic richness of its endosymbiont (Figure 3D) (*24*) led us to explore a potential co-dependency to enable the nutritional specialisation of *O. frankpressi*. We combined highly sensitive profile hidden Markov Models sequence similarity searches with PANTHER annotations to overcome the fast rate of molecular evolution of the *Osedax* lineage (*32*) and reconstruct the eukaryotic biosynthetic pathways that produce amino acids and vitamins in seven symbiotic and asymbiotic annelids, including *O. frankpressi* (Supplementary Table 13). Vestimentifera and asymbiotic annelids (*Owenia fusiformis* and *C. teleta*) can produce all non-essential and conditional essential amino acids, but *O. frankpressi* cannot synthesise the amino acids proline, serine, and arginine (which are non-essential or conditional for mammals) (Figure 4A). However, the endosymbiont can produce these three amino acids (Figure 4A), and proline/hydroxyproline and glycine are abundant in collagen (*42*), the core organic component of bone tissue. Indeed, only one enzyme of the proline biosynthetic pathway remains (pyrroline-5-carboxylate reductase), which is expressed at similar levels in the roots and the rest of the body, unlike most amino acid biosynthetic enzymes that are enriched in roots (Figure 4B, E). Similarly, the entire pathway to synthesise serine from intermediates of glycolysis is missing in *O. frankpressi* (Figure 4C). However, *O. frankpressi* (as other annelids) has an intact glycine cleavage system (Figure 4F), which, coupled with the lack of serine biosynthesis, would favour the conversion of collagen-derived glycine into serine through serine hydroxymethyltransferase (*43*). The two copies of this enzyme are highly expressed throughout *O. frankpressi* (Figure 4F) and could provide an additional source of serine on top of those offered by the diet and endosymbiont. Therefore, *O. frankpressi* shows genomic-inferred metabolic adaptations to its unique bone-eating diet in its gene complement, which differs from the more intact metabolic repertoire of Vestimentifera and other asymbiotic annelids (*14*).

The catabolism of amino acids produces ammonia, a compound that can be toxic, but which can also serve as a substrate for amino acid biosynthesis by both animals and bacteria. Most aquatic organisms excrete excess ammonia to the water, but some aquatic animals and most air-breathing vertebrates shuttle ammonia into the urea cycle leading to urea production (*44*). *Osedax frankpressi* lacks four of the five enzymes of the urea cycle, and only possess arginase (Figure 4D). Interestingly, the urea cycle is also incomplete in the leech *Poecilobdella granulosa* (*45*), another symbiotic heterotrophic annelid with a protein-rich diet that excretes ammonia as waste product. In *O. frankpressi*, the lack of CPS1 is especially significant because this enzyme is the rate-limiting step that mediates the entry of ammonia into the urea cycle; in fact, CPS1 genetic deficiency in humans leads to episodic toxic ammonia levels in the blood (“hyperammonemia”) (*46*). However, *O. frankpressi* additionally lacks urease, and therefore this enzyme is not available to convert ammonia (and carbon dioxide) into urea thus ensuring elevated internal ammonia levels. The only enzyme present in the urea cycle of *O. frankpressi* is arginase, which catalyzes the interconversion of arginine—which the worm likely obtains from bone-derived collagen and the symbiont (Figure 4A)—into ornithine and urea. Although the urea produced by this pathway can be expected to be negligible for ammonia homeostasis, the ornithine may be utilized for producing putrescine and other polyamines that are essential for multiple cellular functions (*47*). Therefore, the amino acid-rich diet and lack of a urea cycle almost certainly implies chronic hyperammonemia in *Osedax*. This would favor amino acid biosynthesis by both *Osedax* and their endosymbionts; however, further functional experiments are needed to test this scenario.

### *Osedax* exhibits lineage-specific expansions of matrix metalloproteinases

As a core component of vertebrate bones, collagen is poised to be a key nutrient for *Osedax* (*23, 25, 28*) and the bone-associated microbiome (*48*). Accordingly, transcriptomic analyses uncovered numerous metalloproteases expressed in the root tissue of *O. japonicus* (*29*), and our gene family evolutionary analyses showed that genes involved in collagen catabolism and extracellular matrix organisation are expanded in the genome of *O. frankpressi* (Figure 2H; Supplementary Figure 4F). Amongst these expanded families, genes annotated as matrix metalloproteases (MMPs) are the largest fraction (24.3%). To investigate how MMPs diversified in *O. frankpressi*, we extracted the reconstructed gene families and functional annotations of symbiotic and asymbiotic annelids to identify sequences containing a metallopeptidase domain (InterPro accession IPR006026). We then reconstructed a phylogeny of the metallopeptidase genes using maximum likelihood and Bayesian approaches (Figure 5A; Supplementary Figures 6, 7). Our analyses recovered all previously described classes of vertebrate MMPs with high statistical support (bootstrap node support >80%) (Figure 5A, highlighted in green) and discovered eight new highly supported invertebrate-specific classes of MMPs, labelled A to H (Figure 5A, highlighted in blue). In addition, we identified two *Osedax*-specific large clades of MMPs, which we referred to as MMP-Os1 and MMP-Os2 (Figure 5A, highlighted in red). The *Osedax*-specific expansions are more closely related to invertebrate than to vertebrate collagenases, in support of previous enzymatic observations that suggested generic proteolysis rather than a true collagenase activity in *Osedax* worms (*23*). The majority of MMPs belonging to MMP-Os1 (37.5%) had a metallopeptidase domain combined with a C-terminal hemopexin-like repeats (IPR018487), thought to facilitate binding to other components of the extracellular matrix (*49*) (Figure 5B; Supplementary Figure 8). As observed with the 12 MMPs reported in *O. japonicus* (*29*), all but two of the 63 MMPs found in *O. frankpressi* are more highly expressed in root tissue than in the rest of the body (Figure 5C), and at least 43 out of 63 (68.25%) have a signal peptide. This suggests the MMPs are excreted across the root-bone interface—similar to bone-degrading osteoclast cells of vertebrate animals (*50*) —allowing *Osedax* to digest bone-derived collagen extracellularly and absorbs the resulting nutrients through the root epithelium for direct consumption, transport to the endosymbiont for further catabolism (*24, 28, 29*), or both. Therefore, the large expansion of MMPs in an otherwise reduced genome is a unique trait to *Osedax* worms and it may be related to their ability to exploit bones from diverse vertebrates, which have collagens with different amino acid sequences and therefore different protease-cleavage sites.

**Figure 5.**
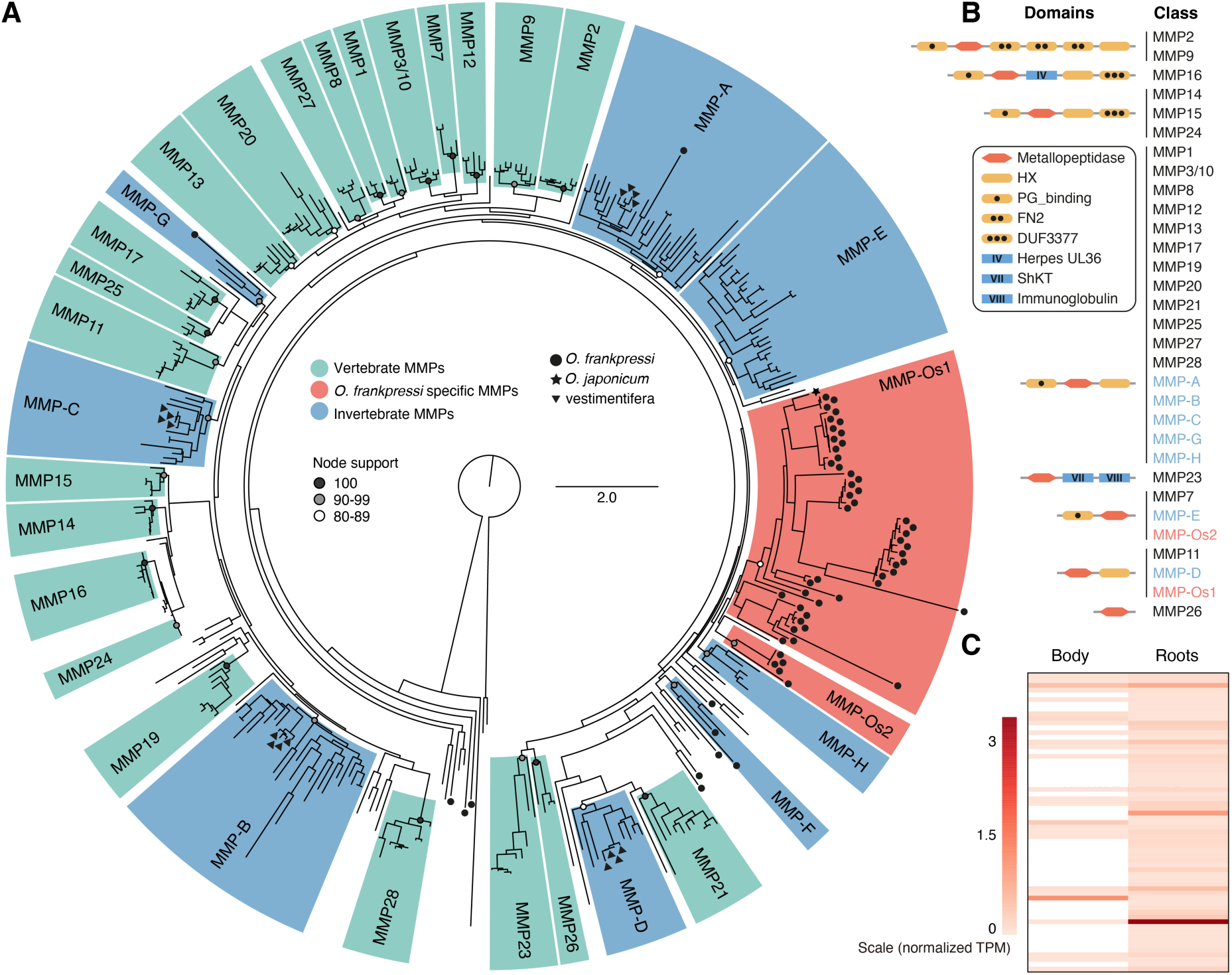
Matrix metalloproteases experienced lineage-specific expansions in *O. frankpressi*. (**A**) Phylogenetic reconstruction of animal matrix metalloproteases (MMPs) based on the metallopeptidase domain. Tree topology is based on maximum likelihood reconstruction and node bootstrap support for each major class is colour coded (white circles show an 80-89 bootstrap support; grey circles indicate a 90-99 bootstrap values and black dots highlight fully supported nodes). Vertebrate-specific MMP classes are highlighted in green and named according to existing literature (*141*). New monophyletic clades of invertebrate MMPs are in blue and named from A to H. *Osedax frankpressi* experienced two independent expansions of MMPs, shown in red and named as MMP-Os1 and MMP-Os2. (**B**) Schematic drawings of the protein domain composition of the different MMP classes recovered in **A**. For each class, only the most abundant domain architecture is shown. A complete characterisation of domain composition of MMPs is in Supplementary Figure 8. Drawings are not to scale. (**C**) Heatmap of normalised expression levels of MMPs in the body and roots of *O. frankpressi*. Most MMPs show higher expression levels in the roots than in the body.

### *Osedax* has a reduced innate immunity repertoire

The establishment of stable and specific host-bacterial associations involves innate immunity genes, which tend to be expanded in Vestimentifera (Supplementary Figure 4E) (*8, 9*) and other symbiotic oligochaetes (*17*). To identify the immune gene repertoire in *O. frankpressi*, we investigated the reconstructed gene families for innate immune pattern recognition receptors corresponding to six major classes, namely lectins, peptidoglycan recognition proteins, Toll-like receptors, scavenger receptors, bactericidal permeability increasing proteins, and NOD-like receptors (*51*). Compared to asymbiotic annelids (i.e., *Owenia fusiformis* and *C. teleta*) and Vestimentifera, *O. frankpressi* has fewer immunity genes in all considered classes (Figure 6; Supplementary Tables 14, 15). This includes a smaller repertoire of Toll-like receptors, which are expanded in Vestimentifera (*8, 9*), and the loss of galectin and a NOD-like receptor, which is a family of cytosolic immune receptors that recognises and trigger inflammatory responses to bacterial pathogens (*52*) that are also largely expanded in Vestimentifera (*9*) (Supplementary Tables 14, 15). Notably, there is no clear association between the expression levels of the different classes of pattern recognition receptors and the body regions and/or tissues of Siboglinidae, yet a C-type lectin is highly expressed in the root tissue of *O. frankpressi* (Figure 6). Together, our findings indicate that *O. frankpressi* and Vestimentifera have significantly different innate immune complements that are simplified in the former and expanded in the latter. This divergence in the repertoire of innate immune gene correlates with the evolution of a novel symbiotic association with *Oceanospirillales* bacteria in *Osedax* worms, and their more dynamic acquisition of endosymbionts compared to Vestimentifera (*40*).

**Figure 6.**
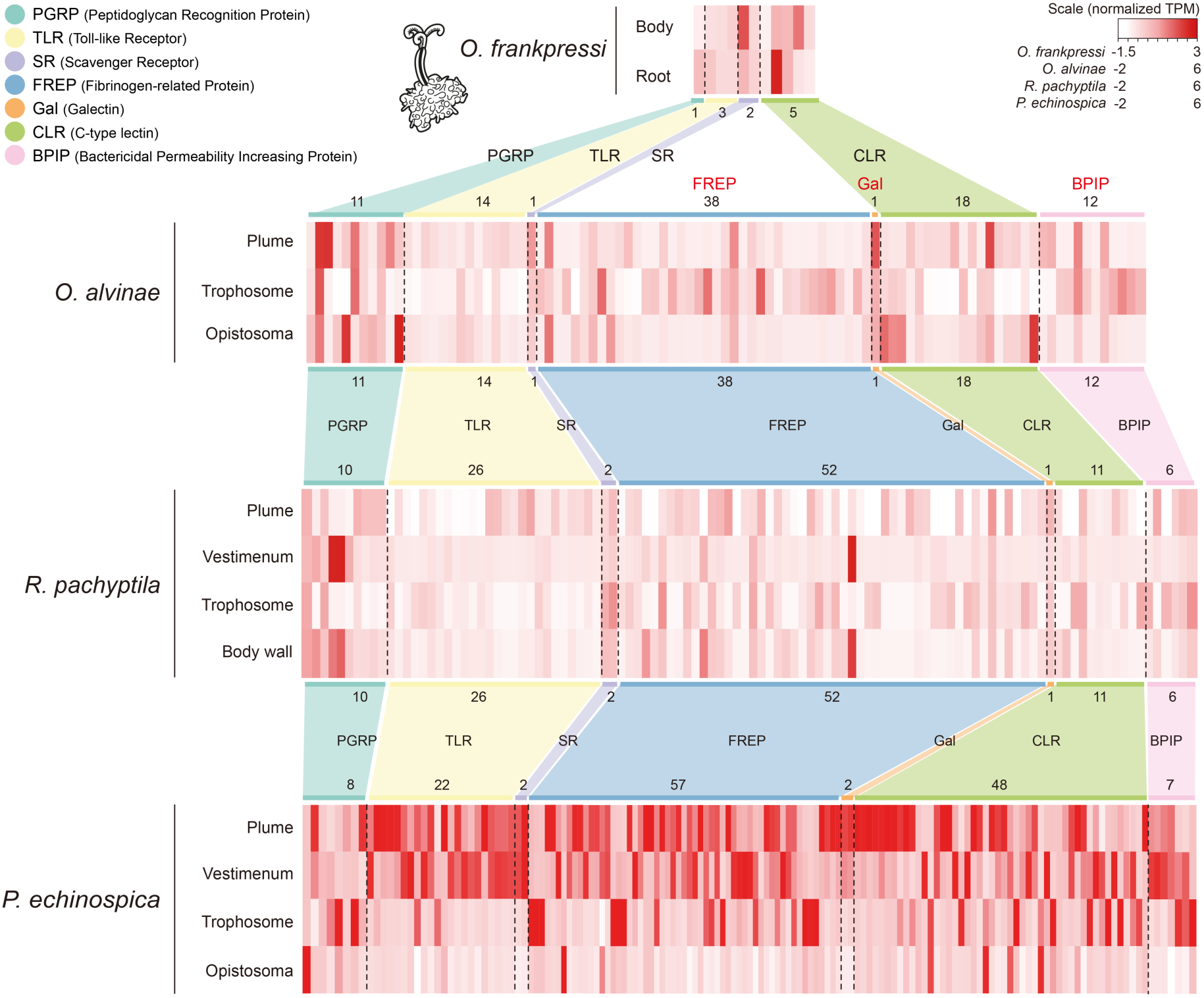
*O. frankpressi* has a reduced innate immune gene repertoire compared to Vestimentifera. Heatmaps of tissue-specific normalised gene expression of innate immune genes in four species of Siboglinidae, including *O. frankpressi* (top) and the Vestimentifera *Oasisia alvinae*, *R. pachyptila* and *P. echinospica*. While Vestimentifera have relatively similar repertoires of innate immune genes, *O. frankpressi* has a much-reduced complement (see Supplementary Tables 14, 15). Notably, innate immune genes do not show a clear tissue-specific expression within or among species of Siboglinidae.

### A conserved developmental toolkit in Siboglinidae

In addition of lacking a gut, Siboglinidae also lacks eyes and any other sensory structure in their most anterior region, the prostomium (*27, 53*). Yet unlike other annelids with unusual body plans such as the leech *H. robusta* (*34*), the genomes of Vestimentifera contain a complete developmental toolkit (*9, 10*). To investigate genes involved in body patterning and organogenesis in the reduced gene set of *O. frankpressi*, we first focused on the repertoire of G protein-coupled receptors (GPCRs). These belong to a large family of evolutionarily related membrane receptors involved in an array of developmental, sensory, and hormonal processes (*54, 55*). We thus compared the genomes of *O. frankpressi*, the Vestimentifera *Oasisia alvinae* and *R. pachyptila*, nine asymbiotic annelids and eight other asymbiotic metazoan lineages for GPCRs, which we then classified by sequence similarity clustering (Figure 7; Supplementary Figure 9; Supplementary Table 16). All siboglinids show a conserved repertoire of GPCRs of class B (secretins), C (metabotropic glutamate receptors) and F (frizzled and smoothened receptors) (Supplementary Figure 9A–C). However, they have a more divergent complement of rhodopsin-like receptors (class A), with three expanded clusters, one of them showing low similarity to leucine rich repeat containing (LRRC) proteins (Figure 7, highlighted in light blue) and the loss of four families, most notably opsins and the Super Conserved Receptor Expressed in Brain (Figure 7, highlighted in light red). Additionally, *O. frankpressi* shows a species-specific expansion of GPCRs (Figure 7; highlighted in light green). Therefore, the lack of photoreceptor GPCRs in both *O. frankpressi* and Vestimentifera suggests an ancestral loss of light perception in Siboglinidae resulting from the colonisation of light-deprived deep marine environments.

**Figure 7.**
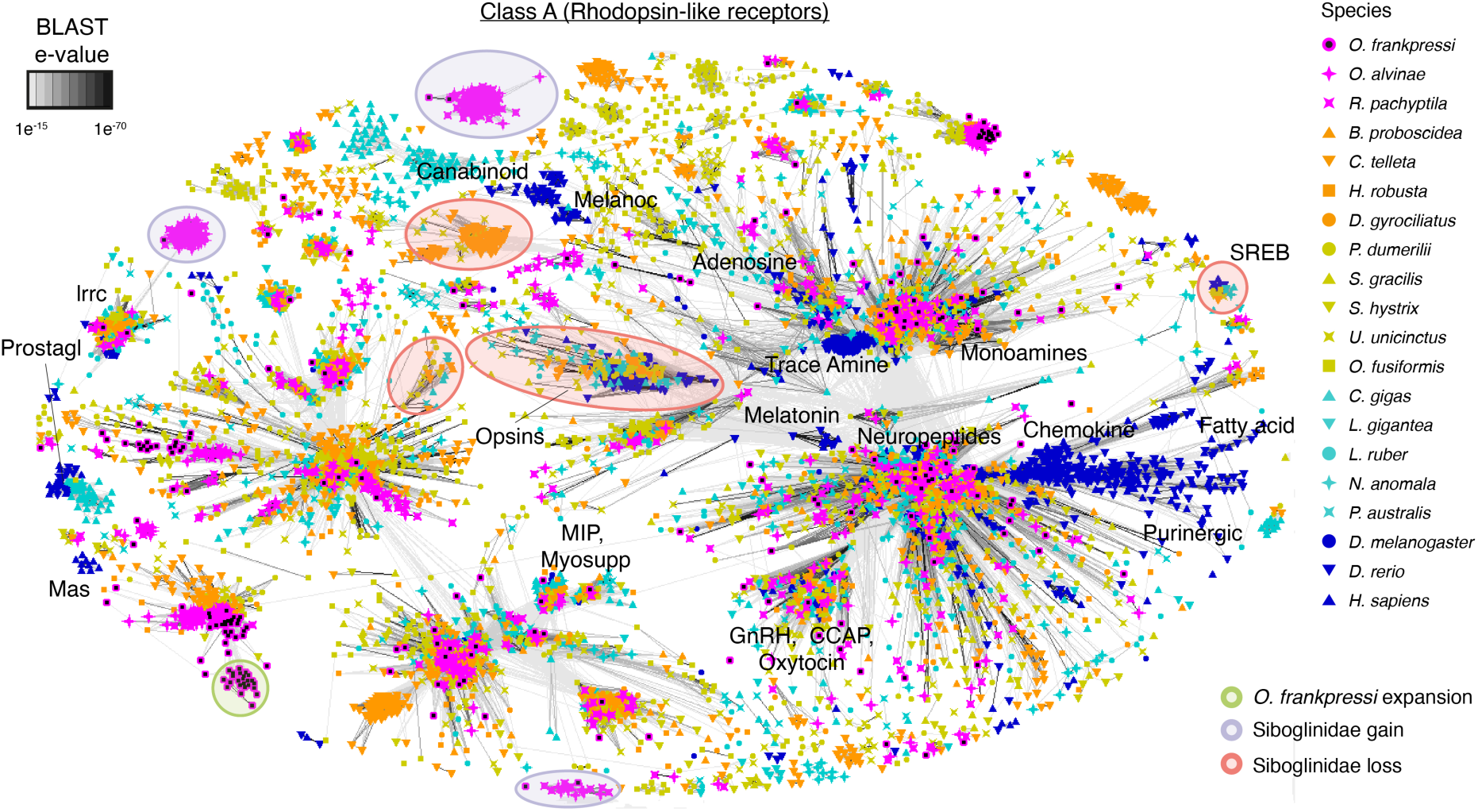
The GPCR complement of Siboglinidae reveals the loss of opsin photoreceptors. Sequence similarity cluster of G-protein couple receptors (GPCRs) of Class A (rhodopsin-like receptors) in Siboglinidae (*O. frankpressi*, *Oasisia alvinae* and *R. pachyptila*; in pink), ten other asymbiotic annelids (in orange and green) and eight other asymbiotic bilaterian lineages (spiralian species in light blue and non-spiralian species in dark blue). Siboglinidae has three clade-specific expansions (highlighted in light purple), one of them weakly related to Leucin-rich containing receptors (lrrc), and *O. frankpressi* has an additional expansion of GPCRs (light green circle in the bottom left). Siboglinidae has also lost four groups of GPCRs present in asymbiotic annelids and other bilaterians, most notably opsins and the Super Conserved Receptor Expressed in Brain (SREB) class.

*Hox* genes are homeodomain-containing transcription factors involved in the anteroposterior regionalisation of bilaterian trunks (*56*) that define a molecular code throughout the many trunk segments in adult Annelida (*33, 57*). Although the bulk of the body of Siboglinidae has only two segments, the posterior end (i.e., the opisthosoma) is often multisegmented, though this is lacking in *Osedax* (*27, 53*). Nevertheless, the *Hox* gene complement in the four studied Vestimentifera is largely conserved, only lacking the gene *Antennapedia (Antp)* (*9, 10*). *Osedax frankpressi* has a similar *Hox* gene repertoire to Vestimentifera, and thus the loss of *Antp* might have also occurred in the last common ancestor of Siboglinidae (Figure 8A; Supplementary Figure 10A). Indeed, the number and complement of transcription factors known to be involved in animal development in *O. frankpressi* are similar to those of Vestimentifera and asymbiotic annelids, except for Basic Leucine Zipper Domain containing proteins (bZIP; PF00170) and zinc finger transcription factors (C2H2-Zn; PF00096), which are reduced (Supplementary Figure 10B), as well as certain specific classes, such as the *ParaHox* genes (Figure 8A; Supplementary Figure 10A). Similarly, *O. frankpressi* retains all major developmental signalling pathways, yet it has a lower number of Notch containing proteins (Figure 8B; Supplementary Figures 10C, 11, 12) and a simplified repertoire of signalling ligands, especially for the Wnt and TGF-β pathways (Figure 8B; Supplementary Figures 10D, 11, 12), as also observed in the annelid with a reduced gene repertoire *D. gyrociliatus* (*32*). Therefore, *O. frankpressi* and Vestimentifera show a similar and generally conserved developmental toolkit, suggesting that changes in gene regulation rather than deviations in the gene complement underpin the development of the divergent adult morphology of Siboglinidae after symbiont acquisition.

**Figure 8.**
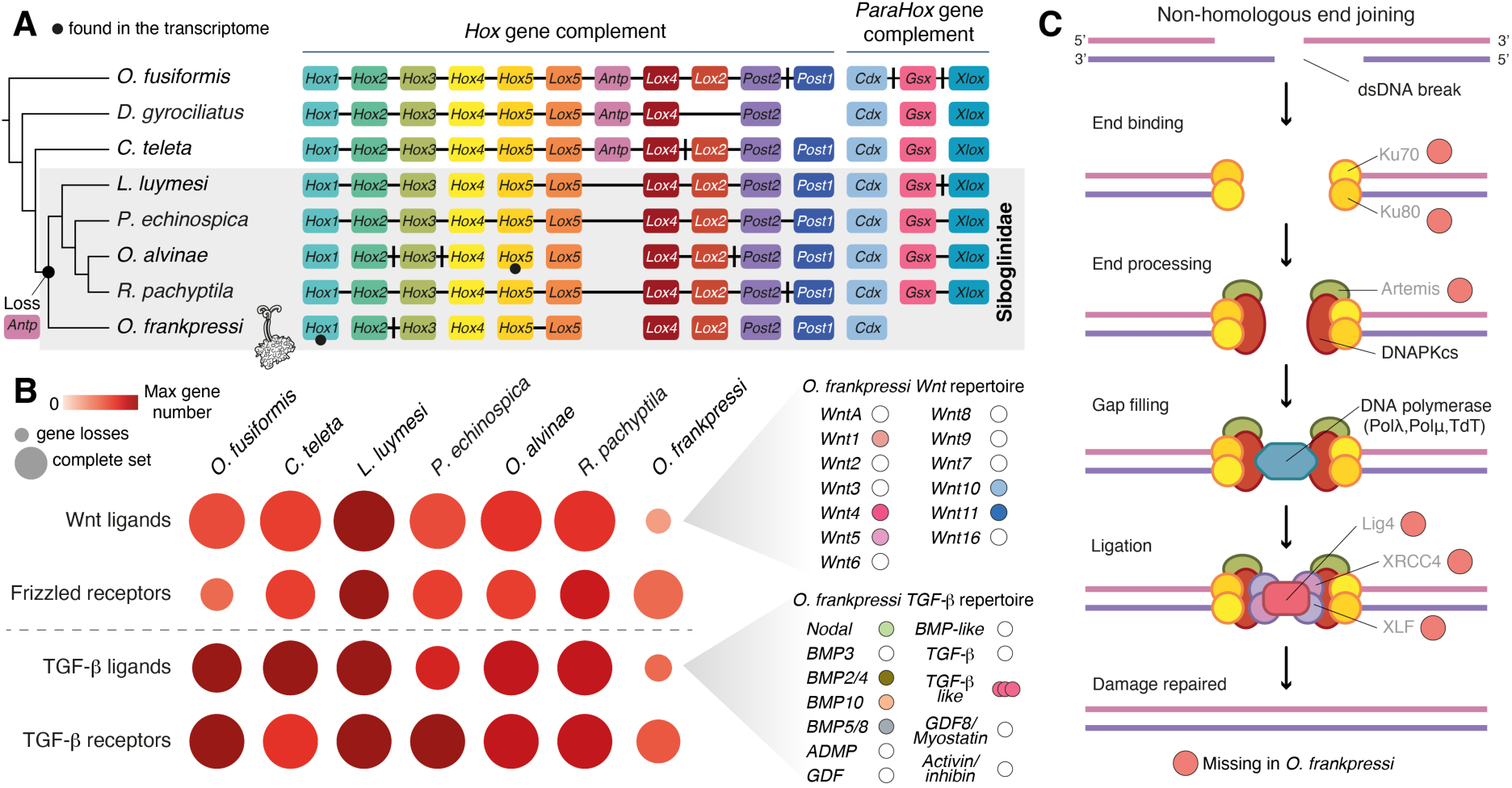
The developmental and DNA repair toolkit of *O. frankpressi*. (**A**) Schematic representation of the *Hox* and *ParaHox* gene complements in Siboglinidae and three other asymbiotic annelid lineages. Each orthologous group is indicated with a different colour, horizontal black lines connecting boxes indicate the genomic linkage, and vertical black lines between boxes indicate that there are interspersed genes between the corresponding *Hox*/*ParaHox* genes. Both *O. frankpressi* and Vestimentifera lack *Antennapedia* (*Antp*), and *O. frankpressi* has in addition lost *Gsx* and *Xlox*. (**B**) Plot summarising the gene number and completeness of the Wnt and TGF-β pathways (based on receptors and ligands) in Siboglinidae and two asymbiotic annelids (*Owenia fusiformis* and *C. teleta*). *Osedax frankpressi* retains the repertoire of receptors but has simplified the complement of Wnt and TGF-β ligands (right side of the panel). (**C**) Schematic representation of the non-homologous end joining DNA repair pathway, highlighting with a red circle the components that are missing in *O. frankpressi*. The lack of this pathway activates the microhomology-mediated end joining repair pathway, which is present in all studied annelids, including *O. frankpressi*, and induces microdeletions that might eventually lead to genome compaction.

### *Osedax* has deficient DNA damage repair mechanisms

Changes in the machinery that repair DNA damage can cause biases in the GC composition of the genome (*58, 59*), and such changes have been associated with genome compaction and gene loss in animals (*60*). The genome of *O. frankpressi* is AT-rich (29.08% GC content versus ∼41% observed in Vestimentifera; Supplementary Figure 2K, M) and genes involved in DNA repair are amongst the families lost in this animal (Supplementary Figure 4D). To investigate how changes in the capacity to repair DNA might correlate with the differences in genome size, gene number, and nucleotide composition among Siboglinidae, we reconstructed all major eukaryotic pathways involved in DNA repair in *O. frankpressi*, Vestimentifera and two slow evolving asymbiotic annelids, using profile hidden Markov Model searches and PANTHER annotations (Supplementary Table 17). Unlike other annelids, *O. frankpressi* has three major DNA repair pathways that are largely incomplete, namely the base excision repair, the non-homologous end joining, and the Fanconi anaemia DNA repair pathway (Figure 8C; Supplementary Figure 13). The base excision repair pathway corrects DNA damage from base lesions caused by deamination, oxidation and methylation, and when impaired, is thought to increase GC to AT base transitions (*61*). Similarly, the lack of the non-homologous end joining pathway—the most common mechanism to repair double-strand DNA breaks (*62*)—triggers the error-prone microhomology-mediated end joining pathway, which causes microdeletions (*63*) and is intact in *O. frankpressi* and all other annelids (Supplementary Figure 13F; Supplementary Table 17). Therefore, the loss of genes involved in the repair of double strand DNA breaks and chemical base modifications might underpin the reduction in genome size and GC content observed in *O. frankpressi* in comparison with Vestimentifera, thus differing from other annelids with reduced genomes, such as *D. gyrociliatus*, whose genome eroded without changes in DNA repair pathways (*32*).

## Conclusions

Our data reveal novel evidence on the genetic interactions and co-dependencies of host and symbiont that allow us to better understand how *Osedax* exploits sunken vertebrate bones as the food source (Figure 9A). Compared to symbiotic Vestimentifera and asymbiotic annelids, *O. frankpressi* shows a fast evolving (*32*), divergent gene repertoire, with gene losses and expansions in key functional groups that support metabolic adaptations to its symbiotic lifestyle (Figure 2, 4A; Supplementary Figure 4D, F). As observed in the marine microbial assemblages on bone surfaces (*48*), the expansion of secreted matrix metalloproteases (Figure 5A) (*29*) combined with the active secretion of acid in the root tissue (*28*) are the most probable mechanisms of bone digestion by the host (Figure 9A). The *Osedax*-microbe association, however, entails a nutritionally unbalanced diet, being deficient in carbohydrates, but enriched in lipids and proline- and glycine-rich proteins (*41, 42*) (Figure 9A). Accordingly, *Osedax*’s *Oceanospirillales* endosymbiont is metabolically versatile (*24, 38*), encoding a significantly larger repertoire of genes involved in sugar and lipid transport as well as carbohydrate, amino acid, and vitamin synthesis compared to the chemolithoautotrophic endosymbionts of Vestimentifera (Figure 3D). An enrichment of secretion systems likely allows the endosymbiont of *Osedax* to outcompete other microbes to occupy the bacteriocytes, as has been shown in other symbiotic systems such as *Vibrio fischeri*-bobtail squid and in the Rhizobia symbionts of legumes (*64, 65*). After the *Osedax-Oceanospirillales* symbiosis is established, the bacteria may transform the bone-derived nutrients acquired by the host into a more diverse set of macronutrients that may be either directly exported to the host or acquired through symbiont digestion (*26*) (Figure 9A). The extensive metabolic repertoire of the symbiont ultimately compensates for the metabolic losses in these worms, while also provisioning the missing nutrients in their specialized diet (Figure 4A–C; Supplementary Figure 4D).

**Figure 9.**
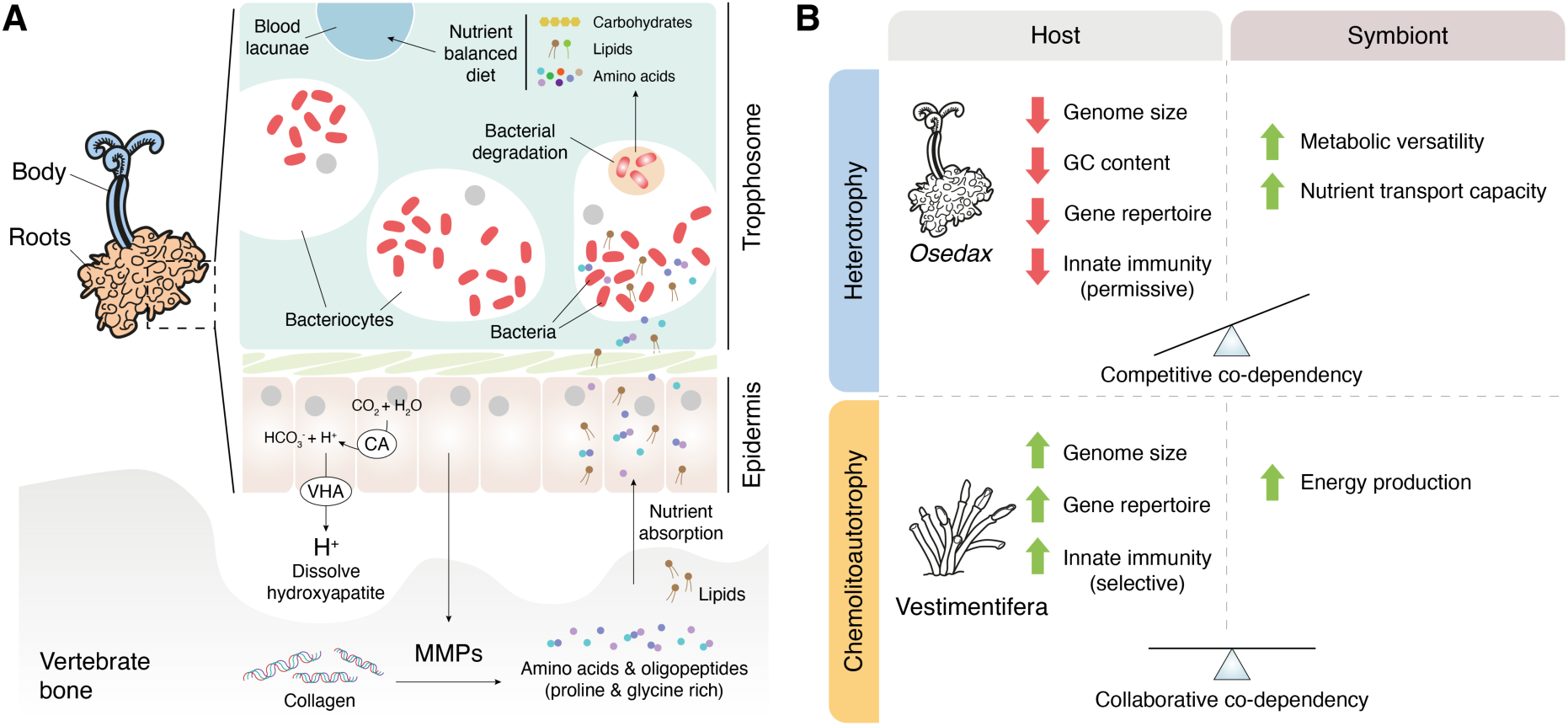
The genomic basis and evolution of heterotrophic symbiosis in *Osedax* worms. (**A**) Schematic drawing of the metabolic interaction for bone digestion between *Osedax* and its endosymbiont (red kidney shaped ovals), which are harboured in the trophosome inside bacteriocytes. The root epidermis secretes acid to dissolve the inorganic component of the bone (via carbonic anhydrase, CA, and V-type H+-ATPase, VHA) and matrix metalloproteases (MMPs) that break collagen, one of the most abundant organic components of the bone, into amino acids and oligopeptides, which are rich in proline and glycine. These amino acids and the lipidic content of the bone are absorbed by the epidermis and used either directly by *Osedax* or transported to bacteriocytes, where they are used by the endosymbiont. The metabolically versatile endosymbiont can transform this unbalanced diet into complex and diverse macronutrients, which are then taken directly or after the digestion of the bacteria by the host. (**B**) *Osedax* and Vestimentifera exhibit markedly different genomic traits. While *Osedax* has a small, AT-rich genome, with many gene losses and a reduced immune repertoire, Vestimentifera tends to show larger genomes, with an enlarged gene complement and rich innate immunity. We hypothesise that the different nutritional relationship between host and symbionts in these two groups might explain, at least partially, these genomic differences. *Osedax* and its endosymbiont co-depend on and compete to exploit the finite, nutritionally unbalanced diet obtained from bones, which might have favoured the evolution of an energetically “cheaper” genome in *Osedax*. In Vestimentifera, however, the endosymbiont acts as a primary producer, which might be able to sustain larger host genomes. Drawings are not to scale.

Symbiotic interactions can impose selective pressures that direct genome evolution—most notably in symbionts(*66*) but also occasionally in hosts (*67*)—triggering changes in genome size (e.g., genome erosion) (*68*), gene content (*69*) and even DNA base composition in favour of AT-rich genomes (*70*). Most of these changes, however, are known for strictly vertically transmitted obligate symbionts of insects. Our study shows that Vestimentifera and *Osedax*, two annelid lineages within Siboglinidae that establish environmentally acquired symbioses, show marked differences in genome structure and composition (Figure 2A–C; Supplementary Figure 2M). While Vestimentifera tends to have larger genome sizes, similar GC content to asymbiotic annelids (*33, 34*), and expanded gene repertoires, *O. frankpressi* has a small, AT-rich genome, with a reduced gene content (Figure 9B). In addition, these Siboglinidae crucially differ in their nutritional symbioses—chemolithoautotrophic in Vestimentifera and heterotrophic in *Osedax*—which enable them as adults to thrive in different ecological niches with different nutritional pressures. In hydrothermal vents and methane seeps, Vestimentifera relies on virtually unlimited inorganic nutrients that are exploited by the endosymbionts, which in their role as primary producers sustain long-lasting collaborative co-dependencies with their hosts (*3, 5*). Decaying bones are, however, nutritionally finite, and thus *Osedax* and their symbiont establish a competitive co-dependency to exploit those nutritionally unbalanced resources (Figure 9B). But while *Oceanospirillales* must retain a diverge metabolic repertoire to accommodate both its free-living and host-bound lifestyles, *Osedax* survives on a homogenous bone-derived diet. We hypothesise that this competitive co-dependency between *Osedax* and its endosymbiont might in turn favour the genomic streamlining—at multiple levels, from size to base composition—of the annelid host (Figure 9B), so that it becomes metabolically “cheaper” and can sustain larger endosymbiotic populations for longer periods. Our findings thus suggest that incipient genome erosion can occur in hosts with horizontally acquired symbionts, and that adaptive genome evolution may differ based on the type of nutritional interactions between the host and symbiont. In the future, dissecting the metabolic co-dependencies between Siboglinidae and their symbionts, including the Frenulata and *Sclerolinum*—the other two major lineages within Sibogliniade— will help to disentangle the role of neutral and adaptive selective pressures in the evolution of these fascinating, but still poorly understood, animal symbioses.

## Materials and Methods

### Specimen collections, gDNA extraction and sequencing

Live adult specimens of *O. frankpressi*, *Oasisia alvinae* and *R. pachyptila* were obtained with deep-sea specialised robots off the coasts of California and Mexico (Supplementary Figure 1C, D). Mexican samples were collected under CONAPESCA permit PPFE/DGOPA-200/18. Ultra-high molecular weight genomic DNA (gDNA) was extracted following the Bionano Genomics IrysPrep agar-based, animal tissue protocol (Catalogue # 80002) from an entire *O. frankpressi* adult female, a piece of trunk (including trophosome) of *Oasisia alvinae*, and a piece of vestimentum of *R. pachyptila*. Long read PacBio sequencing and short read Illumina sequencing was performed at the Genome Centre of the University California Berkeley in a PacBio Sequel II and Illumina Novaseq platforms (Supplementary Table 1).

### Transcriptome sequencing

Total RNA from dissected tissues and body parts of *O. frankpressi* (body and roots), *Oasisia alvinae* (crown, opisthosome and throphosome), and *R. pachyptila* (crown and trunk wall) was extracted with a NEB totalRNA Monarch kit and used for standard strand-specific RNA Illumina library prep. Libraries were sequenced to a depth of 40-50 million paired reads of 150 bases length in a NovaSeq platform (Supplementary Table 1).

### Host genome assembly and quality check

PacBio reads were used to generate an initial genome assembly with Canu v.1.8 (*71*) with options ‘batOptions=’-dg 3 -db 3 -dr 1 -ca 500 -cp 50’. Two rounds of polishing using PacBio reads were performed using Pbmm2 v.1.1.0 (https://github.com/PacificBiosciences/pbmm2) and Arrow (pbgcpp v.1.9.0) (*72*). Short genomic Illumina reads were quality filtered with FastQC v.0.11.8 (*73*) and Cutadapt v.2.5 (*74*), mapped to the polished assembly with BWA v.0.7.17 (*75*) and used for final polishing with Pilon v.1.23 (*76*). The polished versions of the genomes of *O. frankpressi*, *Oasisia alvinae* and *R. pachyptila* were used as input to BlobTools v.2.1 (*77*) to identify and remove contigs with high similarity to bacteria. After decontamination, the host genome assembly was de-haploidised with Purge_Dups v.1.0.1 (*78*). Quality check was performed with BUSCO v.3.0.2 (*79*), to estimate gene completeness of the assembly (Supplementary Table 3), QUAST v.5.0.2 (*80*), and KAT v.2.4.2 (*81*) to assess haplotype removal (Supplementary Figure 2B–D) and potential bacterial remnants.

### Genome size estimations

Short Illumina reads were mapped to the reference host genome assembly with BWA v.0.7.17 and KAT v.2.4.2 (*81*) to count and generate a histogram of canonical 21-mers. GenomeScope2 (*82*) was used to estimate the genome size and heterozygosity (Supplementary Figure 2E–G).

### Symbiont genome assembly and annotation

For *O. frankpressi* and *Oasisia alvinae*, we used Kraken2 v.2.1.0 (*83*) and Krakentools v.0.1 (*83*) to isolate long PacBio reads of bacterial origin. After error correction with Canu v.1.8 (*71*), these PacBio reads were assembled using Metaflye v.2.9 (*84*) followed by ten polishing iterations with options “--pacbio-corr --meta --keep-haplotypes --iterations 10” and final polishing with NextPolish v.1.4.0 (*85*). The resulting assemblies were manually inspected using Bandage v.0.9.0 (*86*), binned with MaxBin2 v.2.2.7 (*87*) and quality checked with CheckM v.1.0.8 (*88*) and MetaQuast v.5.2.0 (*89*). Gene annotation was performed with Prokka v.1.14.5 (*90*) with the “—compliant” option and proteins involved in secretion system were identified by scanning for unordered replicons using the curated HMM profiles of TXSscan in MacSyFinder v.2 (*91*). All coding sequences were assigned KO numbers using BlastKOALA v.2.2 (*92*), which were used as input for KEGG Mapper v.5 (*93*) to analyse the metabolic capabilities of each symbiont. The NCBI COG database (*94*) was used to tag functional categories to the annotated genes. Enrichment analyses of functional categories and Gene Ontology terms were performed with GSEA v.4.2.3 (*95*) and OrthoVenn2 v.2 (*96*). GTDB-Tk v.1.6.0 (*97*) was used for whole genome phylogenetic placement and identification of neighbouring available genomes isolated from free living deep sea bacteria. Circos v.0.69-9 (*98*) was used for genome assembly visualisation.

### Annotation of repeats in host genomes

RepeatModeler v.2.0.1 (*99*) and Repbase (*100*) were used to build a *de novo* library of repeats for the host genome of *O. frankpressi*, *Oasisia alvinae* and *R. pachyptila*. The predicted genes of *Owenia fusiformis* (*33*) and DIAMOND v.0.8.22 (*101*) were used to filter out *bona fide* genes in the predicted repeats with a e-value threshold of 1e-10. Subsequently, RepeatMasker v.4.1.0 (*102*) (Supplementary Tables 5–7) and LTR-finder v.1.07 (*103*) were used to identify and annotate repeats, and RepeatCraft (*104*) to generate a consensus annotation that was used to soft-mask the genome assemblies of the three annelid species. To explore the transposable element landscape, we used the online tool TEclass (*105*) to annotate the TEs identified by RepeatModeler and the scripts “calcDivergenceFromAlign.pl” and a custom-modified version of “createRepeatLandscape.pl”, both from RepeatMasker v.4.1.0, to estimate Kimura substitution levels, which were plotted using ggplot2 v.3.3.0 (*106*). Previously published TE landscapes were included for comparisons (*33*).

### Functional annotation of host genomes

Individual RNA-seq Illumina libraries (Supplementary Table 1) were *de novo* assembled with Trinity v.2.9.1 (*107*) after quality trimming with Trimmomatic v.0.35 (*108*). GMAP v.2017.09.30 (*109*) and STAR v.2.7.5a (*110*) were used to map transcripts and quality filtered Illumina reads to the soft-masked genome assemblies of the corresponding species. For *R. pachyptila*, publicly available datasets (SRA accession numbers SRR8949056 to SRR8949077) were also mapped to the soft-masked genome assembly. In addition, gene transfer format (GTF) files from the mapped reads and curated intron junctions were inferred with StringTie v.2.1.2 (*111*) and Portcullis v.1.2.2 (*112*). All RNA-seq based gene evidences were merged with Mikado v.2.Orc2 (*113*), which produced a curated transcriptome-based genome annotation. Full length Mikado transcripts were used to train Augustus v.3.3.3 (*114*), which was then used to generate *ab initio* gene predictions that incorporate the intron hints of Portcullis and the exon hints of Mikado. Additionally, Exonerate v.2.4.0 (*115*) was used to produced spliced alignments of the curated proteomes of *Owenia fusiformis, C. teleta* and *L. luymesi* that were used as further exon hints for Augustus. Finally, the Mikado RNA-seq based gene evidence and the *ab initio* predicted Augustus gene models were merged with PASA v.2.4.1 (*116*). A final, curated gene set was obtained after removing spurious gene models and genes with high similarity to transposable elements. Gene completeness and annotation quality was assessed with BUSCO v.3.0.2 (*79*). Trinotate v.3.2.1 (*117*), PANTHER v.1.0.10 (*118*) and the online tool KAAS (*119*) were used to functionally annotate the curated gene sets.

### Gene family evolutionary analyses

The non-redundant proteomes of *O. frankpressi*, *Oasisia alvinae* and *R. pachyptila* together with 25 high-quality genomes spanning major groups of the animal tree (Supplementary Table 8) were used to construct orthogroups with OrthoFinder v.2.5.2 (*120*) using DIAMOND v.2.0.9 (*101*) with “–-ultra-sensitive” option. The OrthoFinder output and a published Python script (*32*) were used to infer gene family evolutionary dynamics at each node and tip of the tree. Gene Ontology term enrichment analyses for expanded and lost gene families were performed with the R package “TopGO” v.2.42.0 (*121*).

### Reconstruction of host metabolic pathways and developmental gene sets

PANTHER and Pfam annotations obtained through PANTHER v.1.0.10 (*118*) and Trinotate v.3.2.1 (*117*), respectively, were used to assess for the presence of each enzyme involved in the synthesis of amino acids, vitamin Bs, nitrogen metabolism, glycine degradation, matrix metalloproteases, transcription factors and DNA repair pathways in an array of annelid species. Information about each step in a pathway was collected from MetaCyc (*122*), KEGG (123) and PANTHER (*118*) databases. To analyse the tissue-specific expression of candidate genes in *O. frankpressi*, *Oasisia alvinae* and *R. pachyptila*, quality filtered short Illumina reads were pseudo-mapped to the filtered gene models of each species with Kallisto v.0.46.2 (124) to quantify transcript abundances as Transcripts per Kilobase Million (TPM) values. The R libraries ggplot2 v.3.3.0 (*106*) and pheatmap v.1.0.12 (https://cran.r-project.org/web/packages/pheatmap/index.html) were used to plot expression and abundance heatmaps.

### Reconstruction of innate immune repertoires

The OrthoFinder output was used to identify gene families of innate immune pattern recognition receptors of *O. frankpressi*, Vestimetifera and two asymbiotic annelids, *Owenia fusiformis* and *C. teleta*, with the published pattern recognition receptors of Vestimetifera (*9*) as baits (Supplementary Tables 14, 15). PANTHER and Pfam annotations (see above) of the target proteins were further used to remove sequences that were too short or lacked target domains. TPM expression values (see above) and TBtools v.1.042 (*125*) were used to plot gene expression heatmaps.

### Reconstruction of the G protein-coupled receptor (GPCR) repertoire

Transcriptomes of the focal species were downloaded and processed as described elsewhere (*126*). Multiple sequence alignments of rhodopsin type GPCRs (PF00001), secretin type GPCRs (PF00002), glutamate type GPCRs (PF00003) and frizzled type GPCRs (PF01534) were downloaded from the Pfam webpage (https://pfam.xfam.org) and used to create HMM profiles using hmmer-3.1b2 (*127*). HMMer search was performed with an e-value cut-off of 1e-10. The online version of CLANS (https://toolkit.tuebingen.mpg.de/tools/clans) was used for the initial BLAST comparison for the cluster analysis and edges below 1e-10 were removed. The java offline version of CLANS (*128*) was then used for the cluster analysis. P-value for clustering was set to 1e-30 and edges up to 1e-15 were visualized. Single sequences without connections were deleted (using Linkage clustering for identification). The highly vertebrate specific expanded olfactory GPCR type-A receptors were also deleted as these showed no connections and strongly repulsed all other sequences. Gene clusters were annotated according to the presence of characterized sequences of *Drosophila melanogaster, Homo sapiens, Danio rerio* and *Platynereis dumerilii*.

### Orthology assignments

MAFFT (*129*) with default options was used to align candidate sequences to a curated set of proteins that we obtained either from previous studies (*32, 130*) or manually from UniProt (*131*). Conserved protein domains were retained by trimming by hand the alignment in Jalview (*132*) and the resulting sequences were re-aligned in MAFFT with the “L-INS-I” algorithm (*129*). After a final trim to further remove spurious regions with trimAI v.1.4.rev15 (*133*), FastTree v.2.1.10 (*134*) with default options and IQ-Tree v.2.2.0-beta (*135*) (for matrix metalloproteases) using the options “-m MFP -B 1000”, were used to infer orthology relationships. In addition, for the matrix metalloproteases, posterior probabilities were obtained from Bayesian reconstructions in MrBayes v.3.2.7a (*136*), which were performed using as a prior the LG matrix (*137*) with a gamma model (*138*) with 4 categories to describe sites evolution rate. Four runs with eight chains were run for 20,000,000 generations. FigTree v.1.4.4 (https://github.com/rambaut/figtree) and Adobe Illustrator were used to edit the final trees. CD-Search (*139*) with default options and the Conserved Domain Database (CDD) (*140*) were used to annotate protein domains in the predicted matrix metalloproteases.

## Acknowledgements

We thank members of the Martín-Durán and Henry lab for support and discussions, as well as Gustavo A. Ballén, Ferdinand Marlétaz and the core technical staff at the Department of Biology at Queen Mary University of London for their support. This work was funded by a Wellcome Trust Seed Award in Science to JMM-D (213981/Z/18/Z). JWQ, PYQ, and YNS were supported by the Key Special Project for Introduced Talents Team of Southern Marine Science and Engineering Guangdong Laboratory (Guangzhou) (GML2019ZD0409) and the Major Project of Basic and Applied Basic Research of Guangdong Province (2019B030302004). AMC was funded by a Scripps Postdoctoral Fellowship. Collections for this project were enabled by the Monterey Bay Aquarium and Research Institute and the Schmidt Ocean Institute. Many thanks to Chief Scientists Victoria Orphan and Bob Vrijenhoek, the captains and crews of the R/V *Western Flyer* and R/V *Falkor* and the pilots of the ROVs *Tiburon* and *SuBastian* for crucial assistance in specimen collection.

## Author contributions

JMM-D, LH, GR, and GM conceived and designed the study. GR, NR-K and SG collected the samples; GM assembled and annotated all genomes, performed gene family evolution and metabolic complementarity analyses; BP assembled and annotated the symbiont genomes and contributed to metabolic complementarity analyses; YS, PQ and JQ performed analyses on PRR evolution; DT and GT did GPCR evolutionary analyses; FMM-Z performed Bayesian phylogenetic analyses; MT performed genomic extractions; AMC and Martin Tresguerres contributed to host symbiont metabolic analyses; GM, LH, and JMM-D drafted the manuscript and all authors critically read and commented on the manuscript.

## Competing interests

The authors declare no competing interests.

## Data availability

All sequence data associated with this project are available at the European Nucleotide Archive (project PRJEB55047). Additional files are publicly available in a GitHub repository at https://github.com/ChemaMD.

## Code availability statement

No custom code was used in this study.

## Notes

### Competing Interest Statement

The authors have declared no competing interest.

