## Supplementary Figure 1-13 and Supplementary Tables 1, 2, 3, 5-7, 14 for "The hologenome of *Osedax frankpressi* reveals the genetic interplay for the symbiotic digestion of vertebrate bone"

**Index**

- Supplementary Figures 1 – 13
- Supplementary Tables 1, 2, 3, 5–7, 14 (Word format)
- Supplementary Tables 4, 8–13, 15–24 (Excel format)

### Supplementary Figures

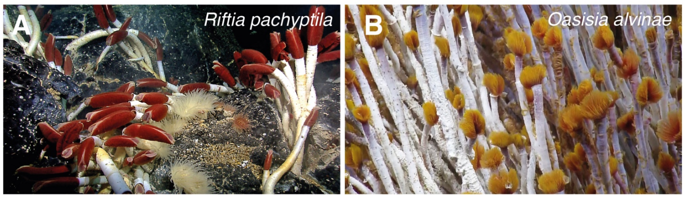


C

| Species | Location | Equipment | Collection date |
|---|---|---|---|
| *Osedax frankpressi* | Whalefall 2893 (Ruby), Monterey Submarine Canyon, California (US) Depth: 2893m | ROV Tiburon, ship R/V Western Flyer | 30 November 2004 |
| *Riftia pachyptila* | Matterhorn vent site, Gulf of California (Mexico) Depth: 3653.5m | ROV SuBastian, ship R/V Falkor | 14 November 2018 |
| *Oasisia alvinae* | Matterhorn vent site, Gulf of California (Mexico) Depth: 3653.5m | ROV SuBastian, ship R/V Falkor | 14 November 2018 |

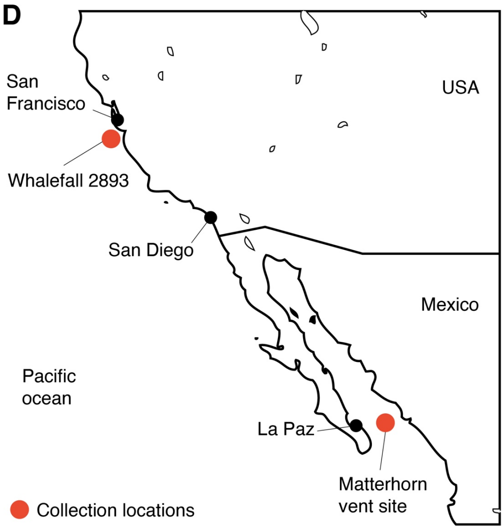


**Supplementary Figure 1. Focal taxa and collection sites.** (**A**, **B**) Photographs of *R. pachyptila* and *Oasisia alvinae* adult worms. Source, wikicommons. (**C**) Table indicating the location, equipment used for the collection, and collection date of the specimens used for genome sequencing for the three species of Siboglinidae herein studied. Voucher specimens are lodged at the Scripps Institution of Oceanography Benthic Invertebrate Collection. (**D**) Schematic map of southern California and northwest Mexico indicating main cities (black dots) and collection sites (red dots).

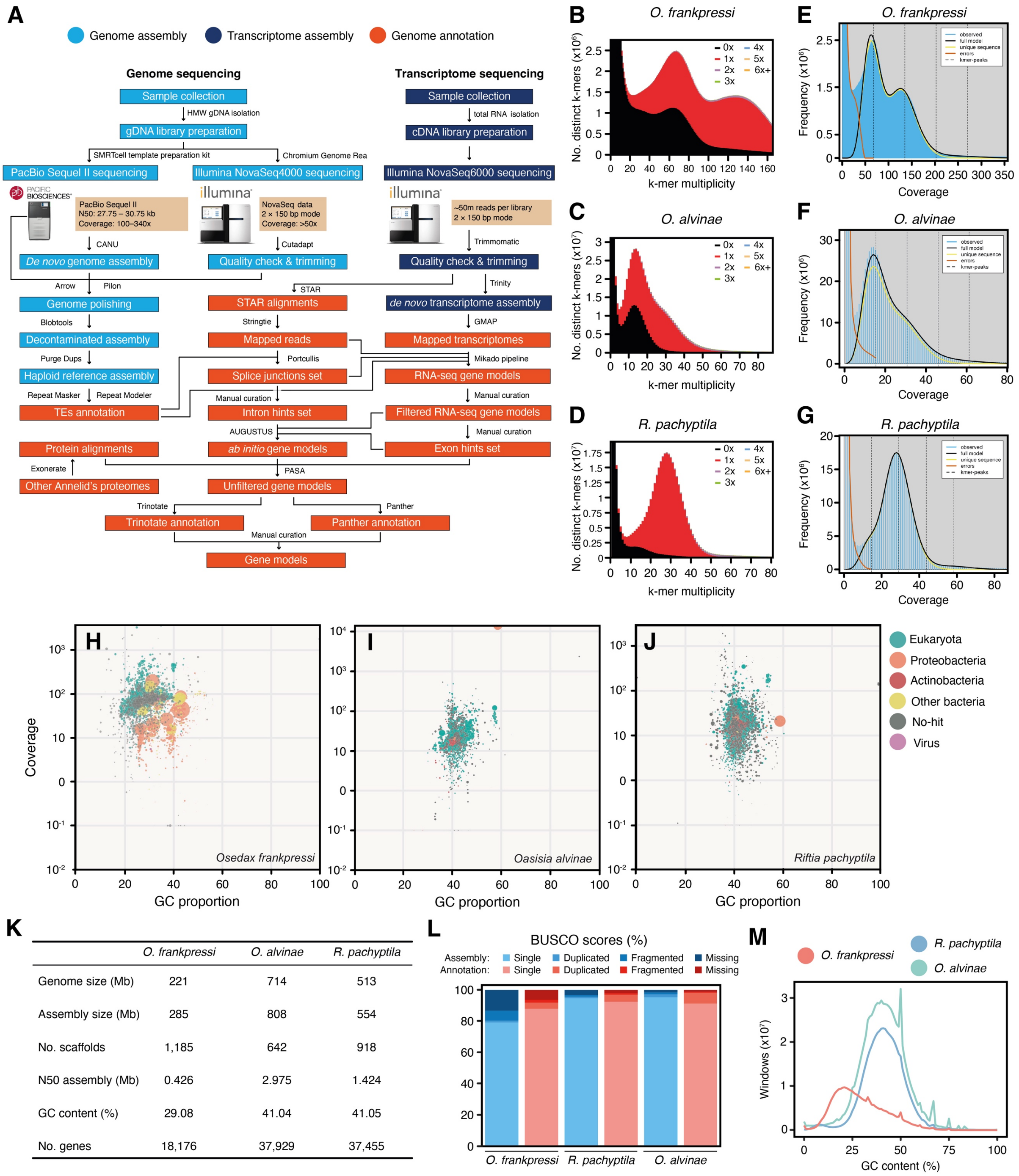


**Supplementary Figure 2. The genomes of *O. frankpressi*, *Oasisia alvinae* and *R. pachyptila*.** (**A**) Schematic diagram outlining the strategy followed to sequence and assemble the genomes (light blue boxes), transcriptomes (dark blue boxes) and functionally annotate the genomes (red boxes) of the three focal species of this study. See Materials and Methods for details. (**B–D**) *k*-mer distribution plots indicating that the genome assemblies for *O. frankpressi*, *Oasisia alvinae* and *R. pachyptila* are largely de-haploidised. (**E–G**)

GenomeScope 2.0 profiles showing *k*-mer based genome size estimations for the three species of study. (**H–J**) Blobtools plots for *O. frankpressi*, *Oasisia alvinae* and *R. pachyptila*, showing eukaryotic contigs in green and prokaryotic light yellow and red. Consistent with the initial biological samples used for genomic extraction (entire animal for *O. frankpressi* and trunk piece including the trophosome for *Oasisia alvinae*), the genomes of those two annelids include their endosymbionts and associated epibiota. (**K**) Table with basic assembly statistics for the three species of study. (**L**) Bar plot indicating the proportions of BUSCO genes in the assemblies (blue bars) and annotations (red bars) of the three focal species. Note how gene annotation improves BUSCO scores in *O. frankpressi*. (**M**) Line plots of GC content in the three Siboglinidae of study.

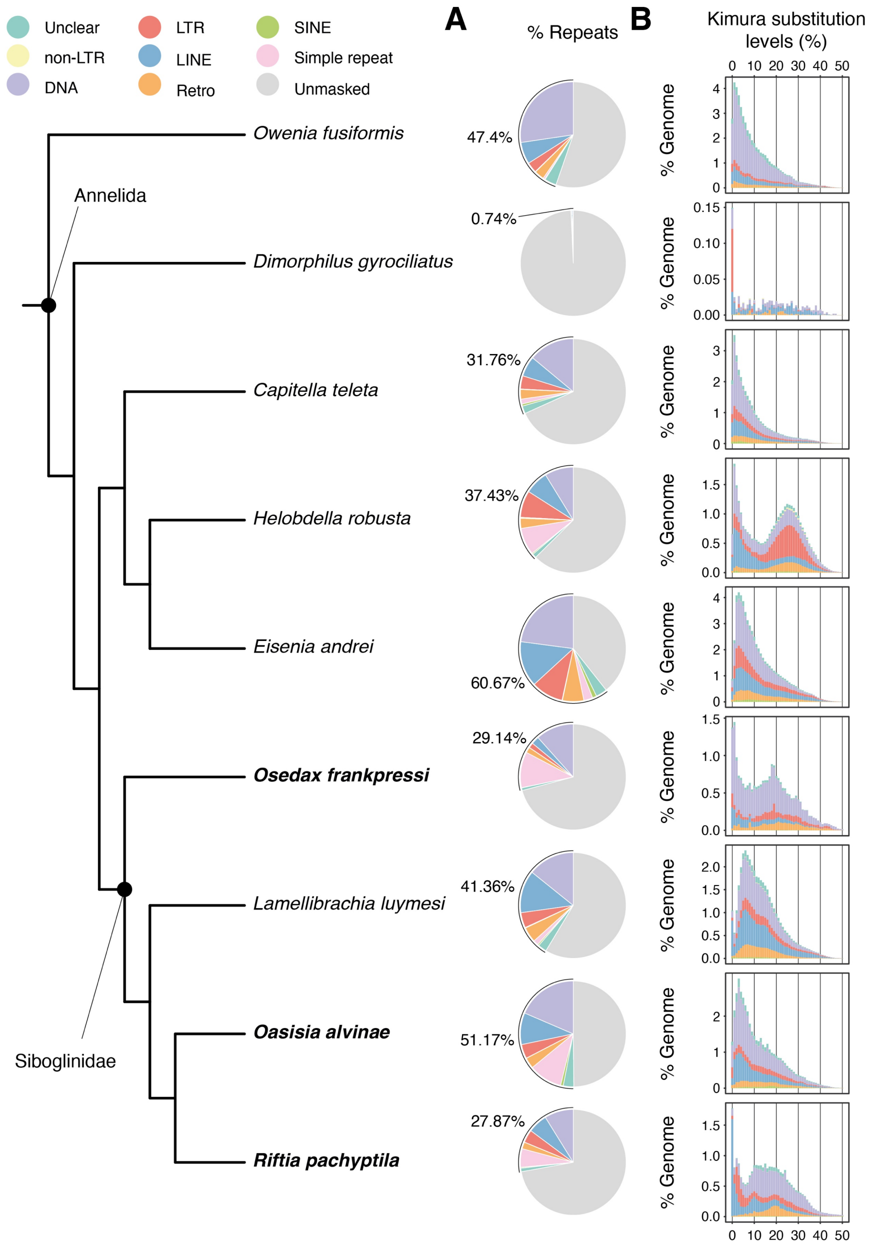


**Supplementary Figure 3. The annelid repetitive landscape.** (**A**) Pie charts showing proportions of repetitive elements in nine annelid genomes under a consensus tree topology (on the left). The genomes of the species sequenced in this study are highlighted in boldface. Despite having a smaller genome, *O. frankpressi* has relatively similar repeat fractions than other Siboglinidae (e.g., *R. pachyptila*) and asymbiotic annelids (e.g., *C. teleta*). (**B**) Distribution plots of Kimura substitution levels of repeat elements and their proportion in the genome for the same nine annelid species indicated on the left of the figure. As observed in other Siboglinidae (e.g., *R. pachyptila*), *O. frankpressi* experienced a past expansion of transposable elements (Kimura substitution levels from 10% to 30%), largely involving DNA transposons.

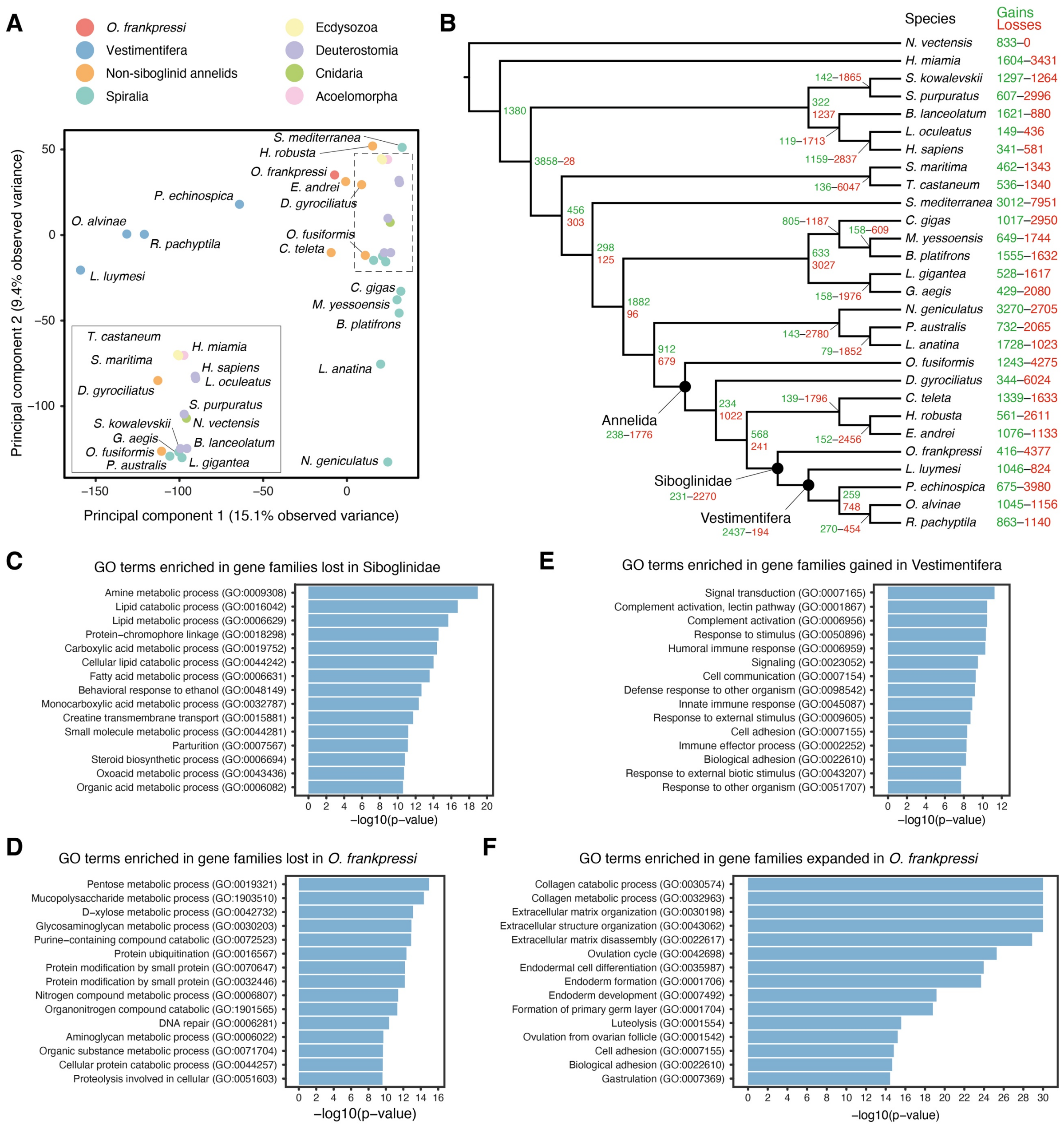


**Supplementary Figure 4. The evolution of the gene repertoire of Siboglinidae.** (**A**) Principal component analysis of 28 highly complete metazoan genomes, including all available Siboglinidae and two symbiotic molluscs (*B. platifrons* and *G. aegis*). While the symbiotic molluscan species are more similar to their asymbiotic relatives, Vestimentifera and *O. frankpressi* are markedly different from asymbiotic annelids with slow rates of molecular evolution, such as *Owenia fusiformis* and *C. teleta*. The area squared by a dashed line in the top right corner is amplified in the bottom left corner to improve taxa identification. (**B**) Patterns of gene family gain (green) and losses (red) for those same 28 metazoan taxa under a consensus tree topology. (**C**–**F**) Bar plots showing the 15 most enriched gene ontology (GO) terms of the Biological Process class in gene families lost in Siboglinidae (**C**) and *O. frankpressi* (**D**), gained in Vestimentifera (**E**), and expanded in *O. frankpressi* (**F**). Most of the losses in Siboglinidae are enriched in GO terms for cellular metabolism, while those gained in Vestimentifera (as per *R. pachyptila*) are enriched in signal transduction, immunity, adhesion, and response to stimulus. *Osedax frankpressi* has lost more gene families enriched in GO terms related to metabolism (mostly carbohydrate) and expanded gene families involved in collagen degradation and extracellular matrix remodelling.

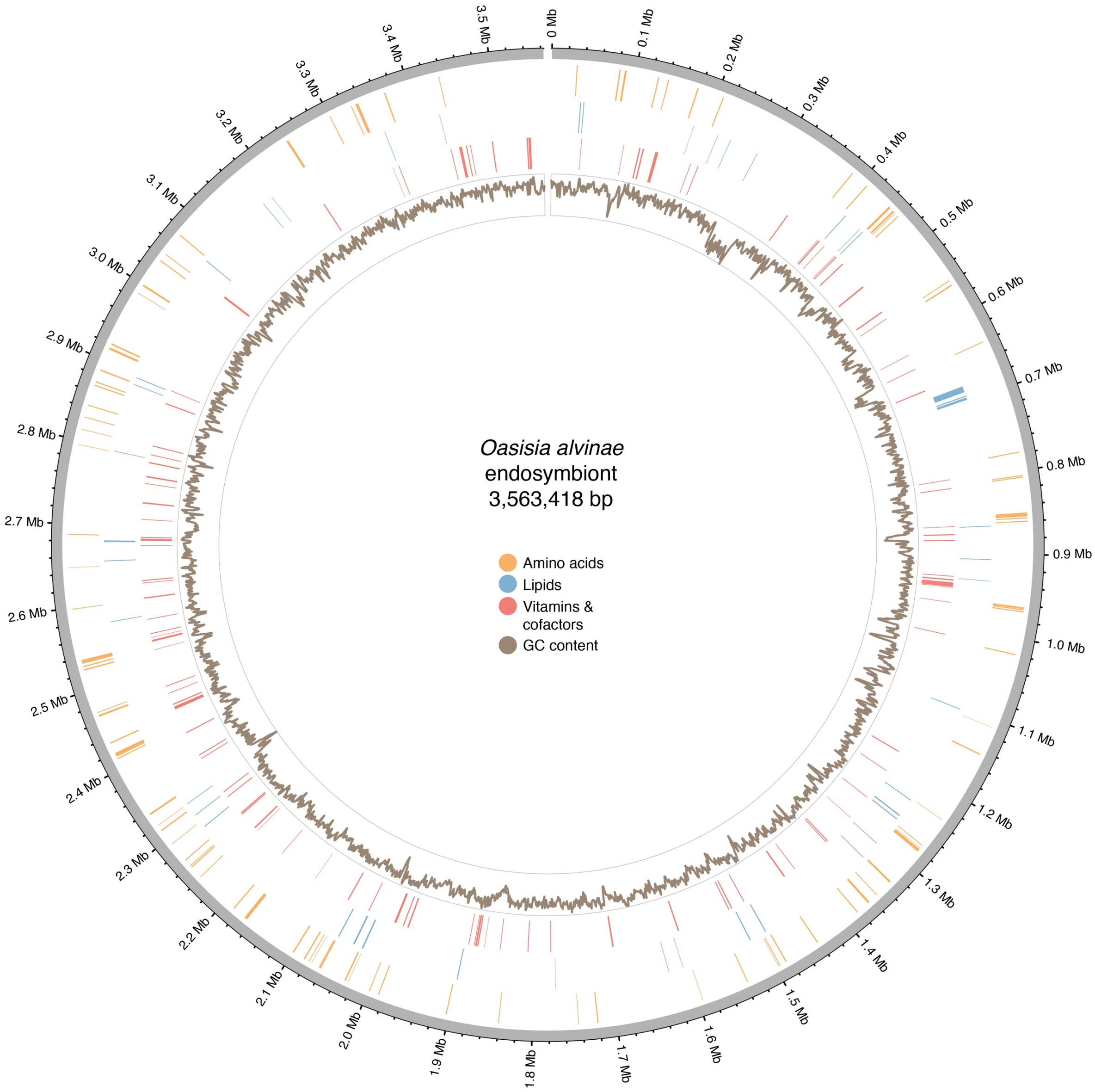


**Supplementary Figure 5. *Oasisia alvinae* endosymbiont.** Circular schematic representation of the genome of *Oasisia alvinae* endosymbiont, assembled into a single contig. The plot shows the genomic location of genes involved in amino acid, lipid, and vitamin/cofactor metabolism (in orange, blue and red, respectively) and the GC content (inner circle; brown colour).

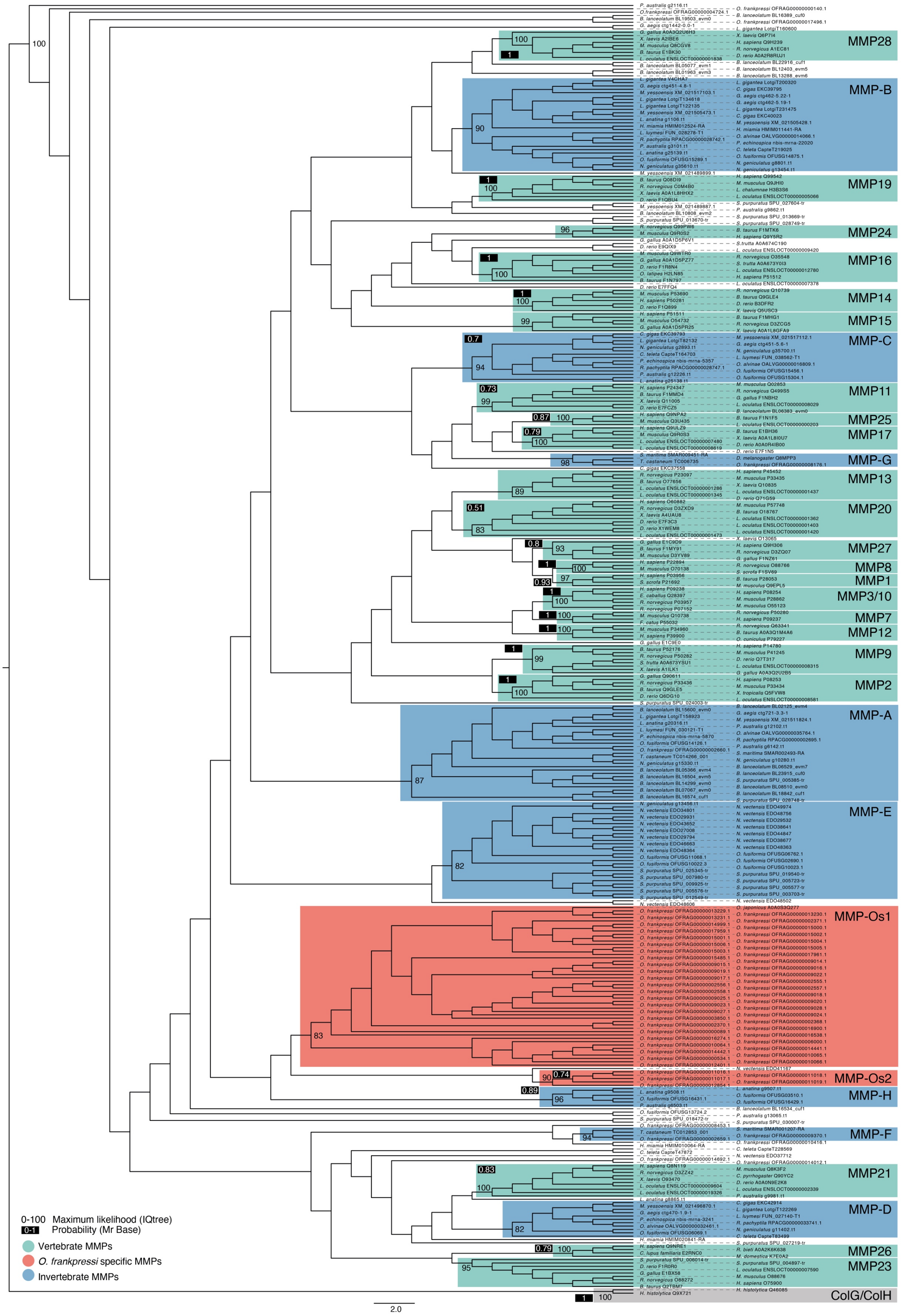


**Supplementary Figure 6. Maximum likelihood phylogenetic reconstruction of animal matrix metalloproteases.** Maximum likelihood consensus tree of animal matrix metalloproteases (MMPs) using the metallopeptidase domain and ColG and ColH bacterial genes as outgroups (grey box). Vertebrate MMPs classes are highlighted by light green boxes, invertebrate MMPs are in blue boxes and the two independent expansions of MMPs in *Osedax* are highlighted with red boxes, showing only bootstrap values for each of the classes.

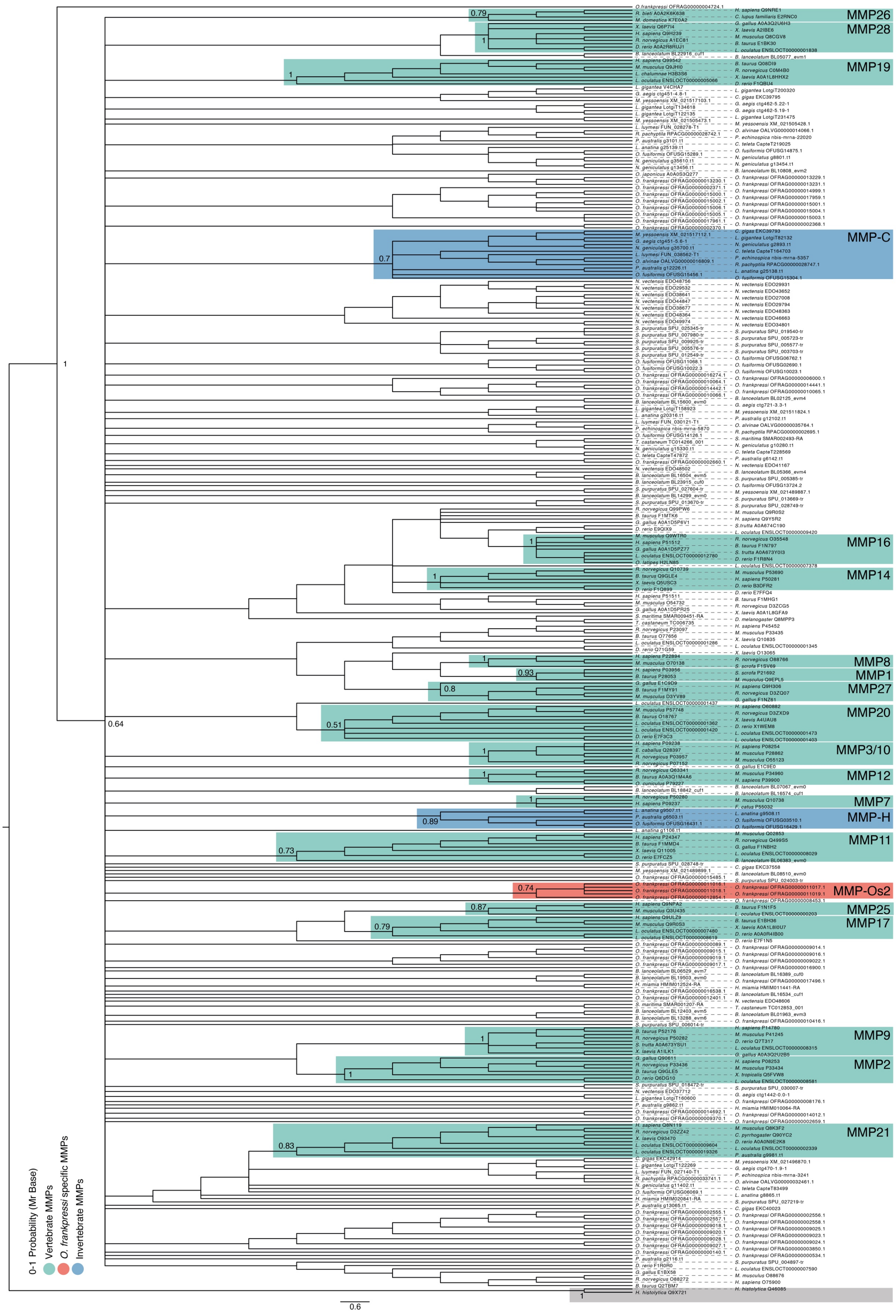


**Supplementary Figure 7. Bayesian phylogenetic reconstruction of animal matrix metalloproteases.** Bayesian consensus tree of animal matrix metalloproteases (MMPs) using the metallopeptidase domain and ColG and ColH bacterial genes as outgroups (grey box). Vertebrate MMPs classes are highlighted by light green boxes, invertebrate MMPs are in blue boxes and the two independent expansions of MMPs in *Osedax* are highlighted with red boxes, showing only posterior probabilities values for each of the classes.

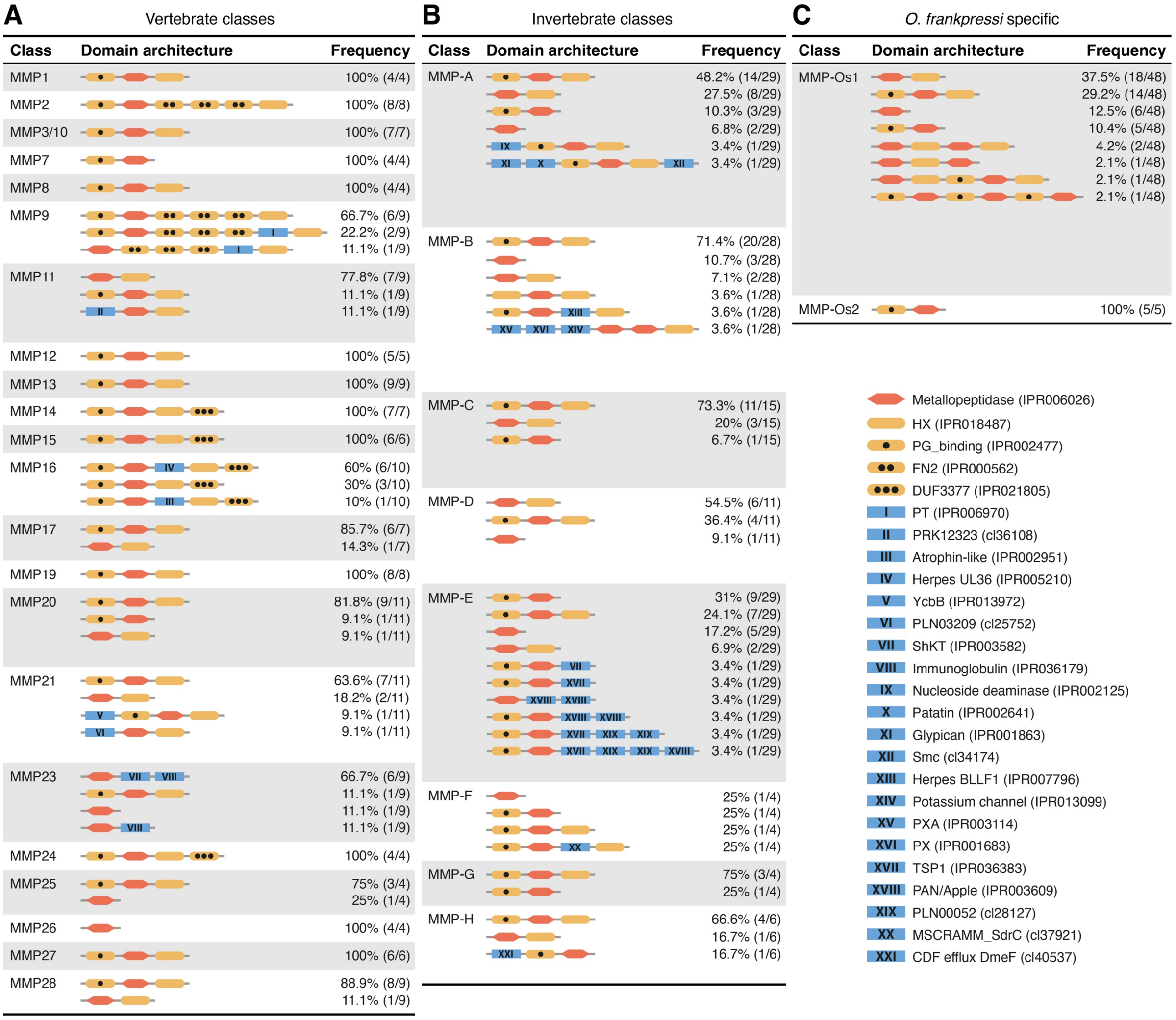


**Supplementary Figure 8. The domain composition of matrix metalloproteases. (A–C)** Schematic representation of the protein domain architecture of matrix metalloproteases in vertebrate (**A**), invertebrate (**B**) and *Osedax* (**C**) classes. Protein domains are coloured based on frequency (in red, the metallopeptidase domain present in all matrix metalloproteases; in yellow domains that are relatively common; and in blue domains that appear in just a reduced number of sequences or classes). The most abundant protein domain configuration for each class is shown in Figure 5B. Drawings are not to scale.

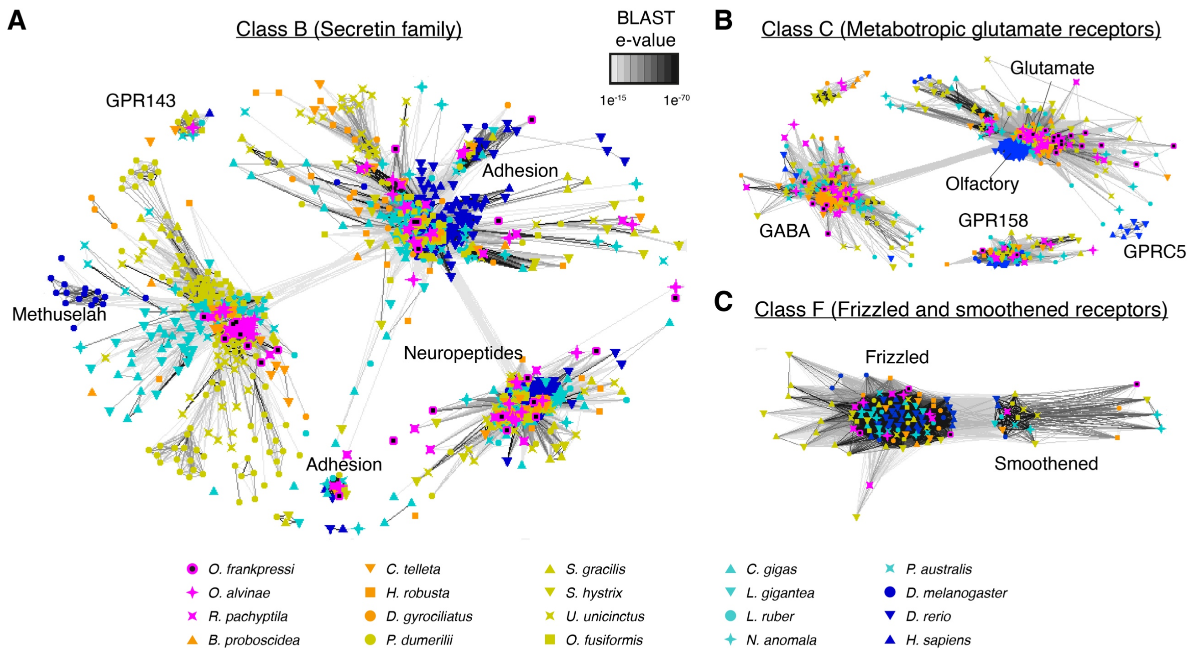


**Supplementary Figure 9. *O. frankpressi* has a normal complement of GPCRs of class B, C and F.** (**A**–**C**) Sequence similarity cluster of G-protein couple receptors (GPCRs) of Class B (secretin family; **A**), class C (metabotropic glutamate receptors; **B**) and class F (frizzled and smoothened receptors; **C**) in Siboglinidae (*O. frankpressi*, *Oasisia alvinae* and *R. pachyptila*; in pink), ten other asymbiotic annelids (in orange and green) and eight other asymbiotic bilaterian lineages (spiralian species in light blue and non-spiralian species in dark blue). All Siboglinidae have representatives of each class and orthogroup within each class.

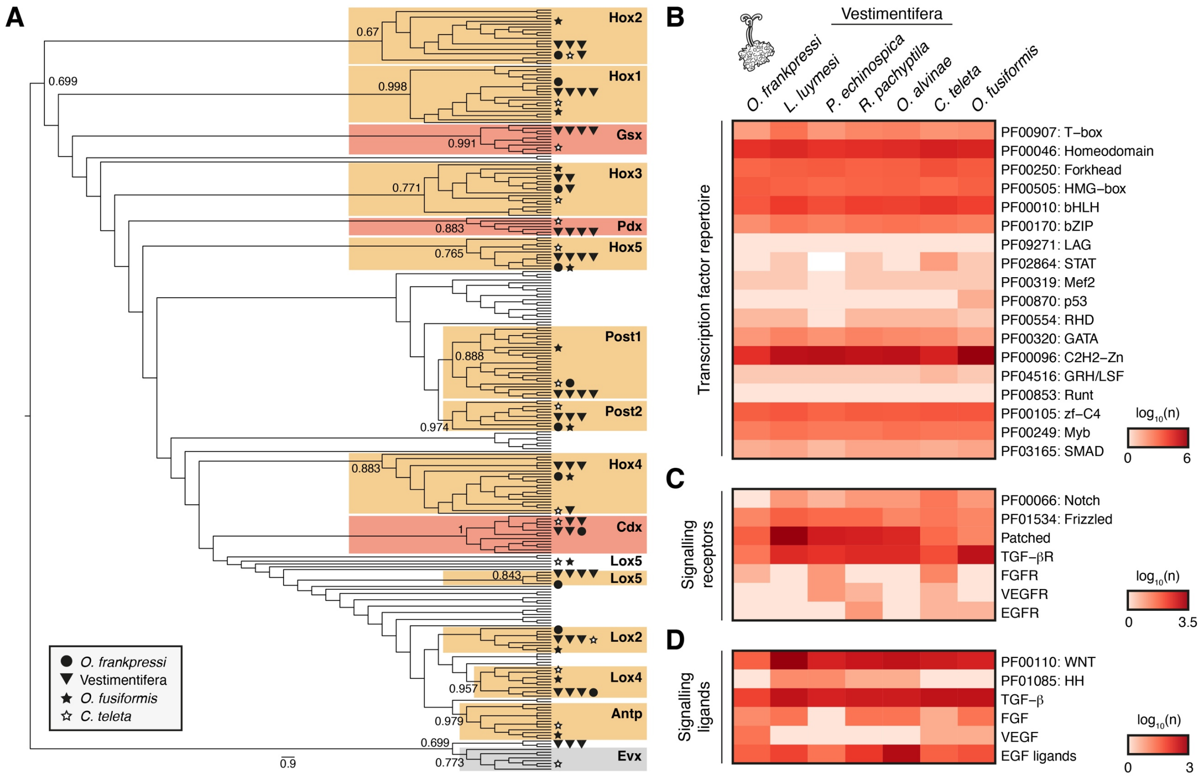


**Supplementary Figure 10. The developmental toolkit and Hox complement in *O. frankpressi*.** (**A**) Orthology inference cladogram of Hox and ParaHox proteins for *O. franpressi* and Vestimentifera, using Evx as outgroup and based on maximum likelihood phylogenetic reconstruction. Numbers show bootstrap support at key nodes of the tree. Symbols indicate the location of annelid sequences in the tree. (**B**–**D**) Heatmaps of the number (in $\log_{10}$ scale) of genes with transcription factor activity (**B**), signalling receptor activity (**C**) and signalling ligands (**D**). The Pfam domain and annotation used for each search is shown on the left. Compared to Vestimentifera and asymbiotic annelids, *O. frankpressi* has a reduced repertoire of Zn finger and bZIP transcription factors, as well as Notch containing receptors. The diversity of Wnt and TGF-b ligands is also lower in *O. frankpressi* (see Figure 8B).

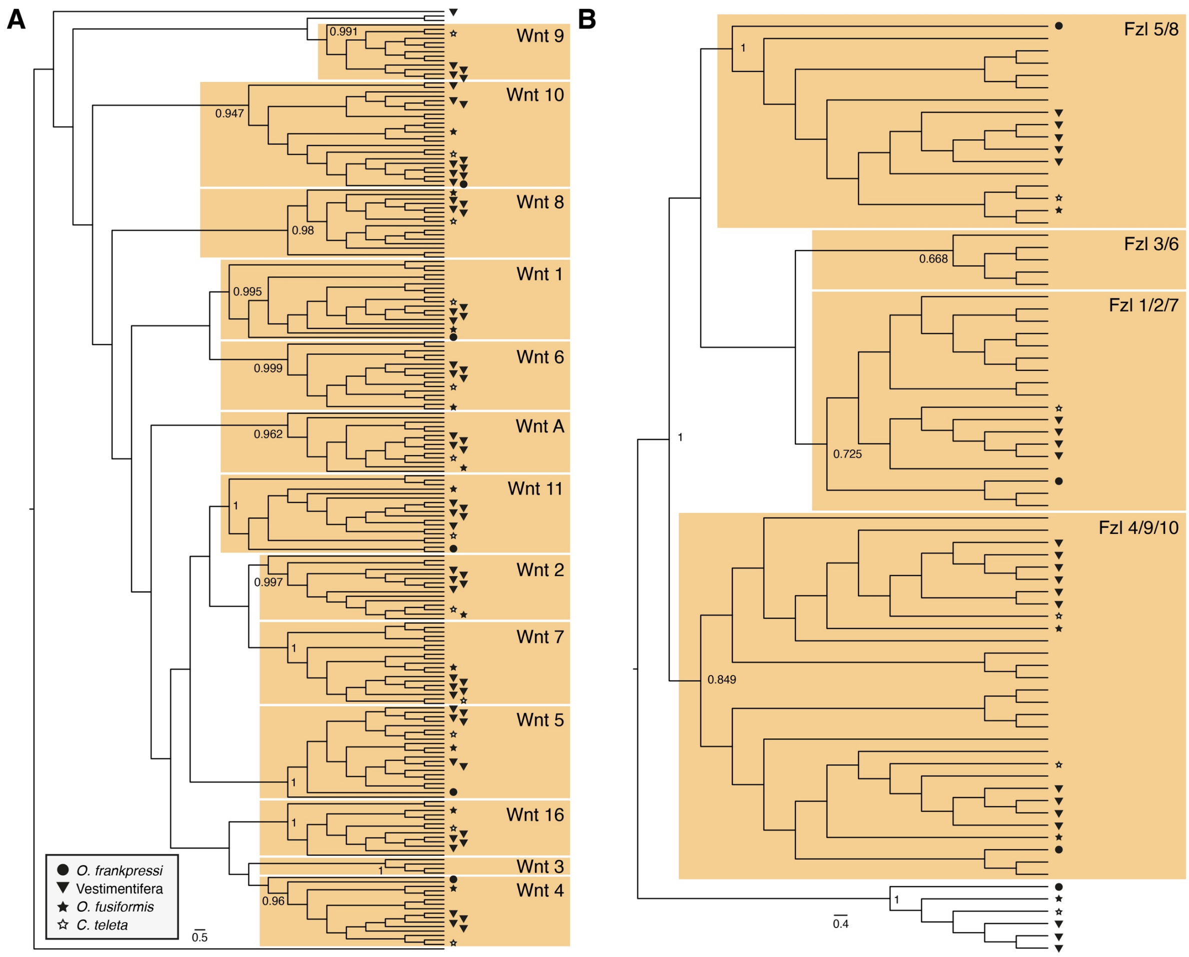


**Supplementary Figure 11. The complement of ligands and receptors of the Wnt pathway in *O. frankpressi*.** (**A**, **B**) Orthology inference cladograms of Wnt ligands (**A**) and receptors (**B**) highlighting each major family with a yellow box, as well as its node support (from 0 to 1). While *O. frankpressi* has an ortholog of each major group of frizzled receptors present in invertebrates (fzl5/8, fzl1/2/7 and fzl4/9/10), it has a reduced diversity of Wnt ligands, with only a copy of Wnt1, Wnt4, Wnt5, Wnt10 and Wnt11. Numbers show bootstrap support at key nodes of the tree. Symbols indicate the location of annelid sequences in the tree.

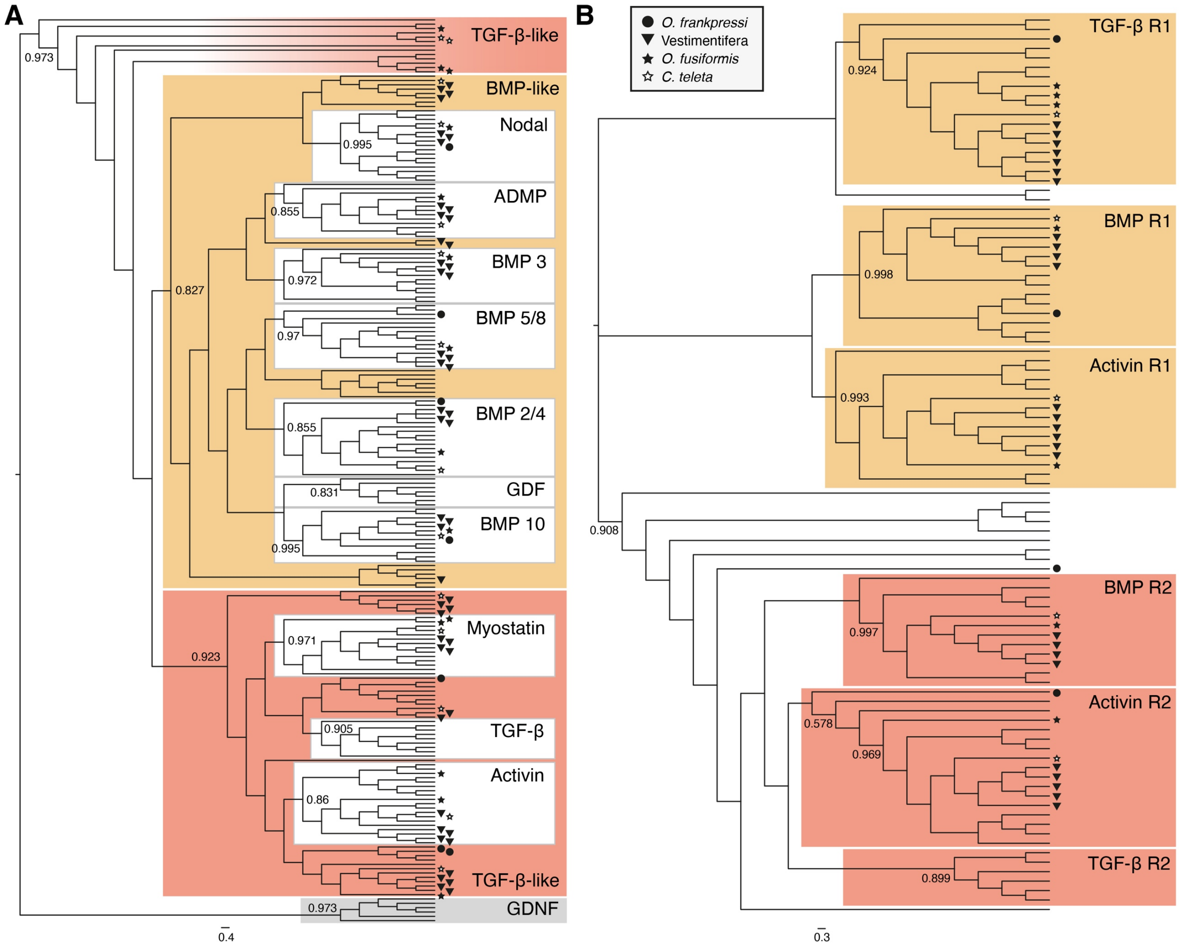


**Supplementary Figure 12. The complement of ligands and receptors of the TGF-b pathway in *O. frankpressi*.** (**A**, **B**) Orthology inference cladograms of TGF-β ligands (**A**) and receptors (**B**) highlighting each major family with coloured boxes (in yellow of the BMP type and in red of the TGF-β type), as well as the node support for each major clade (from 0 to 1). *Osedax frankpressi* has three TGF-β like ligands and a BMP2/4, BMP5/8, BMP10 and Nodal ligands (**A**), and receptors of type 1 and type 2 for BMP and TGF-β like ligands (**B**). Numbers show bootstrap support at key nodes of the tree. Symbols indicate the location of annelid sequences in the tree.

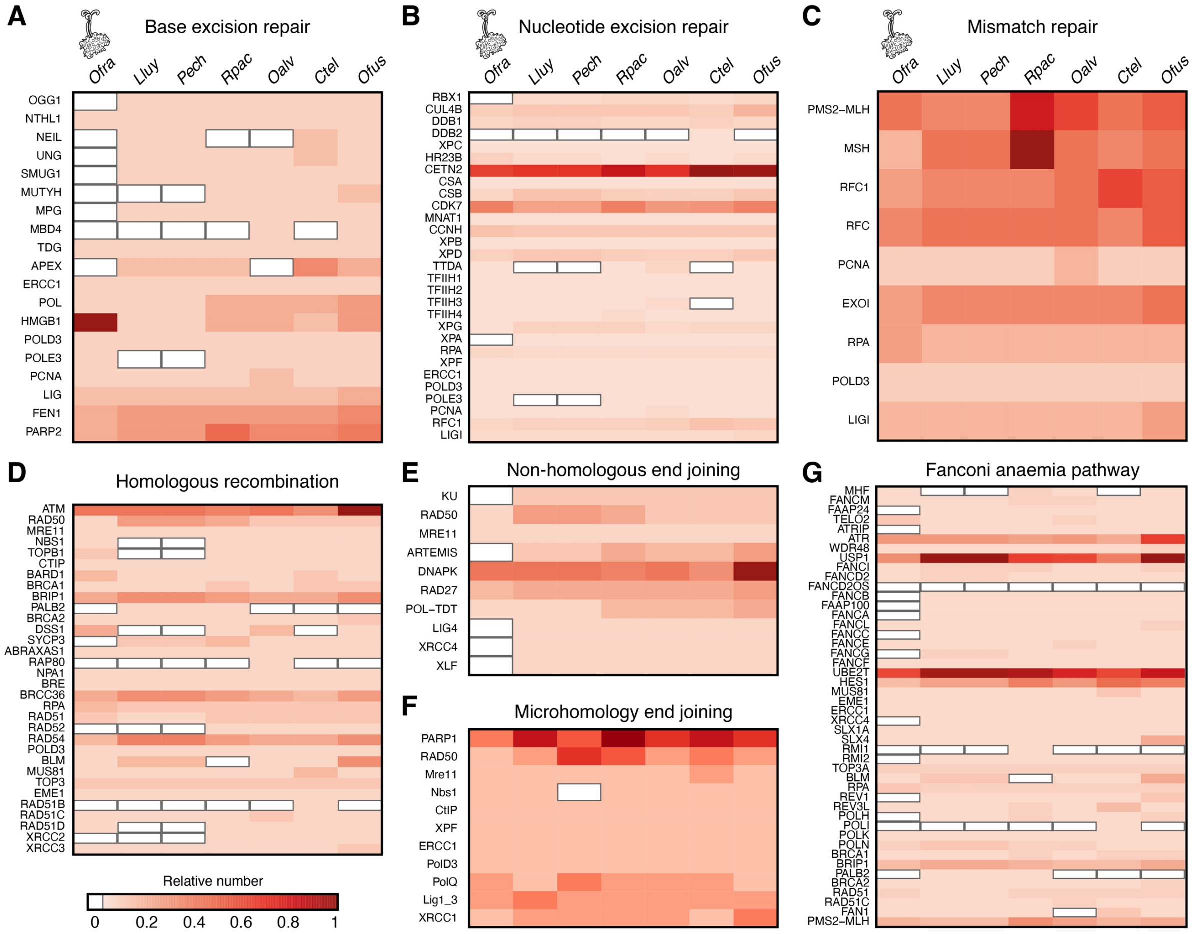


**Supplementary Figure 13. The DNA repair gene complement in *Osedax* and Vestimentifera.** (**A**–**G**) Heatmaps for the presence/absence and number of orthologs for each gene and major pathway involved in the repair of lesions in the DNA based on PANTHER annotations: base excision repair (**A**), nucleotide excision repair (**B**), mismatch repair (**C**) homologous recombination (**D**), non-homologous end joining (**E**), microhomology end joining (**F**) and the Fanconi anaemia pathway (**G**). *Osedax frankpressi* (*Ofra*) shows more gene losses than other Vestimentifera (*L. luymesi*, *Lluy*; *P. echinospica*, *Pech*; *R. pachyptila*, *Rpac*; and *Oasisia alvinae*, *Oalv*) and asymbiotic annelids (*C. teleta*, *Ctel*; and *Owenia fusiformis*, *Ofus*) in the base excision repair, non-homologous end joining and Fanconi anaemia pathways. The potential lack of these fully functioning DNA repair pathways might favour conversion to AT nucleotides and the activation of the microhomology end joining pathway, which can induce microdeletions and genome reduction. Heatmaps show relative values, with 1 representing the maximum number of genes per gene family in the given set of species. Gene loss is indicated by a white rectangle. See Supplementary Table 17 for detailed numbers and exact PANTHER ID numbers.

**Supplementary Tables**

**Supplementary Table 1. List of genomic and transcriptomic resources per species**

| **Species** | **Source** | **Material** | **Platform** | **# reads** |
|---|---|---|---|---|
| *O. frankpressi* | Full body | gDNA | PacBio | 3,282,211 |
| *O. frankpressi* | Full body | gDNA | Illumina | 192,244,931 |
| *O. frankpressi* | Body | Total RNA | Illumina | 129,965,290 |
| *O. frankpressi* | Roots | Total RNA | Illumina | 137,706,423 |
| *Oasisia alvinae* | Trunk | gDNA | PacBio | 5,569,611 |
| *Oasisia alvinae* | Trunk | gDNA | Illumina | 269,878,377 |
| *Oasisia alvinae* | Crown (replicate 1) | Total RNA | Illumina | 41,555,757 |
| *Oasisia alvinae* | Crown (replicate 2) | Total RNA | Illumina | 46,723,618 |
| *Oasisia alvinae* | Opisthosoma (replicate 1) | Total RNA | Illumina | 47,005,676 |
| *Oasisia alvinae* | Opisthosoma (replicate 2) | Total RNA | Illumina | 45,745,399 |
| *Oasisia alvinae* | Trophosome (replicate 1) | Total RNA | Illumina | 42,724,258 |
| *Oasisia alvinae* | Trophosome (replicate 2) | Total RNA | Illumina | 42,151,309 |
| *R. pachyptila* | Vestimentum | gDNA | PacBio | 2,637,279 |
| *R. pachyptila* | Vestimentum | gDNA | Illumina | 78,491,135 |
| *R. pachyptila* | Crown | Total RNA | Illumina | 45,054,210 |
| *R. pachyptila* | Trunk wall | Total RNA | Illumina | 40,456,681 |

**Supplementary Table 2. List of microbial genomes sequenced from *O. frankpressi***

| Group | Bacterium |
|---|---|
| Endosymbiont | Oceanospirillales RS1 copy1<br>Oceanospirillales RS1 copy2 |
| Epibiont | Arcobacter<br>Sulfurospirillum<br>Sulfurimonas copy1 - HighCov<br>Sulfurimonas copy2 - LowCov |
| Free-living | Alphaproteobacteria_rhodospirillales<br>Alphaproteobacteria_kordiimonadales<br>Fusobacteriaceae_Psychrilyobacter<br>Fusibacteraceae_Fusibacter<br>Pseudomonadales_Thioglobaceae<br>Desulfocapsaceae |

**Supplementary Table 3. BUSCO values of the assemblies and the annotations**

| Species | Complete | Single | Duplicated | Fragmented | Missing |
|---|---|---|---|---|---|
| *O. frankpressi* assembly | 80.1% | 79.0% | 1.1% | 6.4% | 13.5% |
| *Oasisia alvinae* assembly | 96.9% | 95.1% | 1.8% | 1.2% | 1.9% |
| *R. pachyptila* assembly | 95.6% | 94.5% | 1.1% | 1% | 3.4% |
| *O. frankpressi* annotation | 91.6% | 90.6% | 1% | 1.9% | 6.5% |
| *Oasisia alvinae* annotation | 97.9% | 96% | 1.9% | 0.7% | 1.4% |
| *R. pachyptila* annotation | 96.8% | 96.4% | 0.4% | 1% | 2.2% |

**Supplementary Table 5. RepeatMasker annotation for *O. frankpressi***

| **Class** | **Subclass** | **Number of elements** | **Length occupied** | **Percentage of sequence** |
|---|---|---|---|---|
| SINEs | | 0 | 0 bp | 0 % |
| | ALUs | 0 | 0 bp | 0 % |
| | MIRs | 0 | 0 bp | 0 % |
| LINEs | | 16554 | 7470495 bp | 2.62 % |
| | LINE1 | 1191 | 262659 bp | 0.09 % |
| | LINE2 | 8930 | 4880920 bp | 1.71 % |
| | L3/CR1 | 102 | 44952 bp | 0.02 % |
| LTR elements | | 5792 | 2589492 bp | 0.91 % |
| | ERVL | 0 | 0 bp | 0 % |
| | ERVL-MaLRs | 0 | 0 bp | 0 % |
| | ERV_classI | 3021 | 728659 bp | 0.26 % |
| | ERV_classII | 427 | 107558 bp | 0.04 % |
| DNA elements | | 58532 | 15266934 bp | 5.36 % |
| | hAT-Charlie | 0 | 0 bp | 0 % |
| | TcMar-Tigger | 2906 | 329535 bp | 0.12 % |
| Unclassified | | 87799 | 26257536 bp | 9.22 % |
| Total interspersed repeats | | / | 51584457 bp | 18.12 % |
| Small RNA | | 320 | 936986 bp | 0.33 % |
| Satellites | | 4999 | 555329 bp | 0.20 % |
| Simple repeats | | 222433 | 29821287 bp | 10.48 % |
| Low complexity | | 17229 | 1377273 bp | 0.48 % |
| **Total bases masked** | | **/** | **82995924 bp** | **29.16 %** |

**Supplementary Table 6. RepeatMasker annotation for *Oasisia alvinae***

| Class | Subclass | Number of elements | Length occupied | Percentage of sequence |
|---|---|---|---|---|
| SINEs | | 31813 | 7301688 bp | 0.90 % |
| | ALUs | 0 | 0 bp | 0 % |
| | MIRs | 31351 | 7254252 bp | 0.90 % |
| LINEs | | 201908 | 70921252 bp | 8.78 % |
| | LINE1 | 6831 | 898275 bp | 0.11 % |
| | LINE2 | 67934 | 23752119 bp | 2.94 % |
| | L3/CR1 | 26088 | 10752788 bp | 1.33 % |
| LTR elements | | 80211 | 15398036 bp | 1.91 % |
| | ERVL | 0 | 0 bp | 0 % |
| | ERVL-MaLRs | 0 | 0 bp | 0 % |
| | ERV_classI | 25807 | 2596412 bp | 0.32 % |
| | ERV_classII | 7459 | 1192793 bp | 0.15 % |
| DNA elements | | 186008 | 56807183 bp | 7.03 % |
| | hAT-Charlie | 1757 | 376153 bp | 0.05 % |
| | TcMar-Tigger | 209 | 30159 bp | 0 % |
| Unclassified | | 623188 | 164893139 bp | 20.41 % |
| Total interspersed repeats | | / | 315321298 bp | 39.03 % |
| Small RNA | | 12830 | 3636450 bp | 0.45 % |
| Satellites | | 27203 | 5678762 bp | 0.70 % |
| Simple repeats | | 362245 | 81655285 bp | 10.11 % |
| Low complexity | | 10824 | 905243 bp | 0.11 % |
| **Total bases masked** | | **/** | **405156532 bp** | **50.15 %** |

**Supplementary Table 7: RepeatMasker annotation for *R. pachyptila***

| Class | Subclass | Number of elements | Length occupied | Percentage of sequence |
|---|---|---|---|---|
| SINEs | | 21320 | 3026230 bp | 0.55 % |
| | ALUs | 0 | 0 bp | 0 % |
| | MIRs | 21320 | 3026230 bp | 0.55 % |
| LINEs | | 87276 | 43599677 bp | 7.88 % |
| | LINE1 | 11208 | 1624069 bp | 0.29 % |
| | LINE2 | 27602 | 8280880 bp | 1.50 % |
| | L3/CR1 | 29494 | 28314275 bp | 5.11 % |
| LTR elements | | 90618 | 12941344 bp | 2.34 % |
| | ERVL | 0 | 0 bp | 0 % |
| | ERVL-MaLRs | 0 | 0 bp | 0 % |
| | ERV_classI | 10399 | 881879 bp | 0.16 % |
| | ERV_classII | 1250 | 318412 bp | 0.06 % |
| DNA elements | | 79305 | 12117760 bp | 2.19 % |
| | hAT-Charlie | 1058 | 164379 bp | 0.03 % |
| | TcMar-Tigger | 0 | 0 bp | 0 % |
| Unclassified | | 195756 | 45050193 bp | 8.14 % |
| Total interspersed repeats | | / | 116735204 bp | 21.09 % |
| Small RNA | | 809 | 524256 bp | 0.09 % |
| Satellites | | 34434 | 9644285 bp | 1.74 % |
| Simple repeats | | 192588 | 30818651 bp | 5.57 % |
| Low complexity | | 6510 | 578920 bp | 0.10 % |
| **Total bases masked** | | **/** | **154286816 bp** | **27.87 %** |

**Supplementary Table 14: Pattern recognition receptors in selected annelids[1]**

| PRRs | Ofra | Oalv | Rpac | Pech | Lluy | Ofus | Ctel |
|---|---|---|---|---|---|---|---|
| Lectins | | | | | | | |
| C-type lectin (CTLs) | 8 | 26 | 14 | 47 | 77 | 85 | 52 |
| Fibrinogen-related protein (FREPs) | 5 | 62 | 62 | 66 | 80 | 92 | 82 |
| Galectin | 0 | 1 | 1 | 2 | 1 | 6 | 4 |
| Peptidoglycan recognition protein (PGRP) | 1 | 14 | 10 | 9 | 18 | 19 | 7 |
| Scavenger receptor (SR) | 2 | 5 | 3 | 3 | 3 | 5 | 10 |
| Toll-like receptor (TLR) | 9 | 50 | 35 | 23 | 46 | 75 | 21 |
| Bactericidal permeability increasing protein (BPIP) | 1 | 14 | 6 | 7 | 8 | 2 | 8 |
| Nod-like receptor (NLR) | 0 | 60 | 16 | 20 | 23 | 0 | 1 |

[1]Species abbreviations: Ofra, *O. frankpressi*; Oalv, *Oasisia alvinae*; Rpac, *R. pachyptila*; Pech, *P. echinospica*; Lluy, *L. luymesi*; Ofus, *Owenia fusiformis*; Ctel, *C. teleta*.
